# Gene recruitments and dismissals in argonaut octopus genome provide insights to pelagic lifestyle adaptation and shell-like eggcase reacquisition

**DOI:** 10.1101/2021.11.08.467834

**Authors:** Masa-aki Yoshida, Kazuki Hirota, Junichi Imoto, Miki Okuno, Hiroyuki Tanaka, Rei Kajitani, Atsushi Toyoda, Takehiko Itoh, Kazuho Ikeo, Takenori Sasaki, Davin H. E. Setiamarga

## Abstract

The paper nautilus, *Argonauta argo*, also known as the greater argonaut, is a species of octopods distinctly characterized by its pelagic lifestyle and by the presence of a spiral-shaped shell-like eggcase in females. The eggcase functions by protecting the eggs laid inside it, and by building and keeping air intakes for buoyancy. To reveal the genomic background of the species’ adaptation to pelagic lifestyle and the acquisition of its shell-like eggcase, we sequenced the draft genome sequence of the species. The genome size was 1.1 Gb, which is the smallest among the cephalopods known to date, with the top 215 scaffolds (average length 5,064,479 bp) covering 81% (1.09 Gb) of the total assembly. A total of 26,433 protein-coding genes were predicted from 16,802 assembled scaffolds. From these, we identified nearly intact HOX, Parahox, Wnt clusters and some gene clusters probably related to the pelagic lifestyle, such as *reflectin, tyrosinase*, and *opsin*. For example, *opsin* might have undergone an extensive duplication in order to adapt to the pelagic lifestyle, as opposed to other octopuses, which are mostly the benthic. Our gene models also discovered several genes homologous to those related to calcified shell formation in Conchiferan Mollusks, such as Pif-like, SOD, and TRX. Interestingly, comparative genomics analysis revealed that the homologous genes for such genes were also found in the genome of the octopus, which does not have a shell, as well as the basal cephalopods *Nautilus*. Therefore, the draft genome sequence of *A. argo* we presented here had not only helped us to gain further insights into the genetic background of the dynamic recruitment and dismissal of genes for the formation of an important, converging extended phenotypic structure such as the shell and the shell-like eggcase, but also the evolution of lifestyles in Cephalopods and the octopods, from benthic to pelagic.

## Introduction

The paper nautilus, or the argonaut *Argonauta argo* is a member of Argonautoidea, a superfamily of octopods (Cephalopoda, Octopodiformes), but has specialized characters not found in other octopus species. It is a cosmopolitan species distributed in the global tropical and subtropical open seas (Norman, 2000). Phylogenetic analyses have placed *A. argo* together with its congener (e.g. *A. hians*), forming a monophyletic Argonautidae, and then, form a sister relationship with the blanket octopuses (e.g. *Tremoctopus*), and thus further forming the superfamily Argonautoidea (Hirota et al., 2021; Strugnell et al., 2006; Sanchez et al., 2018). Distinct synapomorphies of this superfamily, which could also be found in *A. argo*, are the extreme female-biased sexual size dimorphism, a comparatively large and entirely transformed hectocotylus that is coiled in a pouch below the eye until maturity, and the transferring of spermatophores to the female mantle cavity by hectocotylus detachment (Naef, 1923; Bello, 2012). Although the consensus phylogeny also suggested that Argonautoidea split from benthic ancestral octopods, members of the superfamily including *A. argo* are fully adapted to the holopelagic lifestyle and thus does not need to have any contact with the seafloor throughout its lifecycle. Several studies have suggested that the holopelagic lifestyle was probably achieved by evolutionary acquisitions of distinct characters enabling members of Argonautoidea to keep afloat in midwater and to egg brooding away from the sea floor (cf. Naef 1923; Packard & Wurtz 1994; Young 1985; Bizikov 2004). Extreme adaptations of this group to their midwater habitat have masked their evolutionary origins. Buoyancy in argonauts was probably obtained after their ancestors had already become pelagic, potentially via the pelagic paralarval or juvenile stages found in many benthic octopuses with small eggs (Finn and Norman 2010).

One conspicuous character separating Argonautidae, a family which includes all argonauts (genus *Argonauta*), with the rest of Argonautoidea is the presence of a biomineralized eggcase in females, which external morphology mimics the spirally-wound shells of *Nautilus* and the extinct ammonites (Scales, 2015; Stevens et al., 2015). The eggcase is thought to protect the eggs laid inside or, as well as taking in air for maintaining buoyancy (Finn and Norman 2010). As such, the re-acquisition of this shell-like structure was probably important, because it helps *Argonauta* to maintain its holopelagic lifestyle. Previous observations have maintained that the “shell-like” eggcase is not a “true” shell (the Conchiferan shell) (Naef, 1923).

The evolutionary story of shell formation and loss in Cephalopods is interesting in itself. Although being classified as a member of the Conchifera, a subphylum of Mollusks composed of members with external shells biomineralized with calcium carbonate, except for the basally diverged Nautiloids (Setiamarga et al., 2021a), extant Cephalopods mostly degenerated their shells, resulting in the complete shell loss in octopods, and vestigial shells in some decapods (squids and cuttlefishes) (Kröger et al., 2011). True Conchiferan shells are formed through the secretion of proteins from the mantle tissue, made from aragonite and calcite, have the nacreous layer and intricate microstructures (Jackson et al., 2009; Kocot et al., 2016; Jackson et al., 2017), which has evolved since at least in late Ordovician (Vendrasco et al., 2011; 2013). An extant member of the basal Cephalopods, the Nautiloid *Nautilus pompilius* apparently also forms their shells this way (Marie et al., 2009; Setiamarga et al., 2021a). However, despite convergence in their general external morphology, the eggcase of *Argonauta* is not considered as a true Conchiferan shell but an evolutionary innovation of the genus (Naef, 1923; Scales, 2015). It is formed through the secretion of related proteins from their arms (Naef, 1923; Scales, 2015) and has different biomineralization and microstructural profiles (Revelle and Fairbridge 1957; Mitchell *et al*. 1994; Nixon and Young 2003; Saul and Stadum 2005; Oudot *et al*. 2020). Previously, we conducted an extensive multi-omics analysis on the eggcase of two argonaut species, *A. argo* and *A. hians*, which samples were obtained from the Sea of Japan (Setiamarga et al., 2021b). Two important points relevant to our present study could be taken from our previous one: (1) almost no Conchiferan homologous SMP, including those of the basal Cephalopoda *Nautilus pompilius* (Setiamarga et al., 2021a) was present in the eggcase matrix of the two argonaut species, and (2) Conchiferan SMP homologs (or homologous domains) were also found in the genome of the shell-less octopods, *Octopus bimaculoides*. These points thus indicate that our result was in agreement with the result of morphological observations, which maintains that the eggcase is not a homologous structure of the shell. However, the observations have also caused other questions: Are the SMP genes not used in the eggcase formation still retained in the genomes of the argonauts? Comparative genome analyses across Cephalopoda, and among different representative species of Conchifera, are thus needed to answer this question. Such genomic level comparative studies would also give important insights on the evolution of holopelagic lifestyle at the genetic level.

Until very recently, the lack of genome data had prevented us from understanding the genetic basis of Cephalopod biology, and even molluscan biology. This was remedied by recent reports of various Cephalopod genomes, such as the genomes of *O. bimaculoides* (Albertin et al., 2019), *Euprymna scolopes* (Belcaid et al., 2019), *Architeuthis dux* (da Forsa et al. 2020), and the basal Cephalopod *Nautilus pompilius* (Zhang et al., 2021; Huang et al., 2021). Comparative genomics studies of these genomes have allowed us to identify notable characteristics of Cephalopod genomes except *Nautilus*, such as: (1) the average genome size of around 3 Gigabases (Gb), which is slightly bigger than that of other molluscan species (Gregory 2021), (2) highly rearranged genome with transposable element expansion, which have caused the genomes to be highly repetitive in nature (Albertin et al., 2015; da Fonseca et al., 2020), (3) lineage specific duplication of certain types of genes (Yoshida et al., 2011), and (4) whole transcript-wide adenosine to inosine (A-to-I) RNA editing (Alon et al. 2015; Liscovitch-Brauer et al. 2017). These genomic characteristics have thus suggested that the coleoid Cephalopods have intriguingly different genomes from “standard” metazoan genomes. Another interesting point is that such differences were apparently evolutionarily acquired in ancestral Coleoids, which members show similar body plans and morphology in general (Young 1971) despite their ancient divergences (Decapodiformes (squid and cuttlefishes) vs. Octopodiformes (vampire squid and octopuses) = 242 ± 38 million years ago (Mya); Nautiloidea vs. Coleoidea = 415 ± 60 Mya) (Kröger et al., 2011; Vinther et al., 2012; Sanchez et al., 2016). Therefore, additional genomic data, especially of the Octopodiformes, will allow us to trace the ancestral chromosomes of Cephalopods and their transition within Mollusks, which in the end might help to unravel the evolutionary origin of these “genomic idiosyncrasies” These are major obstacles to tracing the ancestral chromosomes of cephalopods and their transition within the Mollusks.

Here, we report a high-quality draft genome assembly of the greater paper nautilus / greater argonaut *Argonauta argo*. We found that this species has an exceptionally small genome size, making the species an ideal species for genomic studies. Although studies targeting argonauts have not progressed because it is difficult to keep in aquaria, we have access to the location in the Sea of Japan, where fresh and living samples of this species could be easily obtained by fixed nets from June to August (Sakurai and Kono, 2010). Using obtained genome data, we focusedly discussed the evolution of some interesting genomic features such as those related to shell evolution, eggcase formation, and color vision, in order to gain insights to the genomic basis of the adaptation to the open-ocean holopelagic lifestyle of this species in particular, and the evolution of Cephalopods in general.

## Results and Discussion

### The draft genome assembly of *A. argo*

We generated a draft genome from a single individual of a female argonaut obtained from a fixed net set on the coasts of Oki Island, Shimane Prefecture, Japan (Figure 1). A total of 1.34 Gb was assembled from input sequences obtained from genome sequencing with 201× coverage (107× PE, 24× 3 kb MP, 24× 6 kb MP, 24× 10kb MP, and 24× 15kb MP). 57,036 scaffolds of various lengths were assembled, with the top 215 scaffolds larger than 1000 kb (average length 5,064,479 bp), covering 81% (1.09 Gb) of the total assembly. Half of assembled scaffolds (N50) were of 6.18 Megabases (Mb) or longer, reflecting high contiguity. These statistics (Table S1) thus showed that our *A. argo* draft genome sequence ranks among the top quality draft genomes of Molluscs, and the most comprehensive for Cephalopods. For example, our N50 indicates that our *A. argo* draft genome is twice as long as that of the Hawaiian bobtail squid *E. scolopes* (Belcaid et al. 2019).

**Figure 1.**
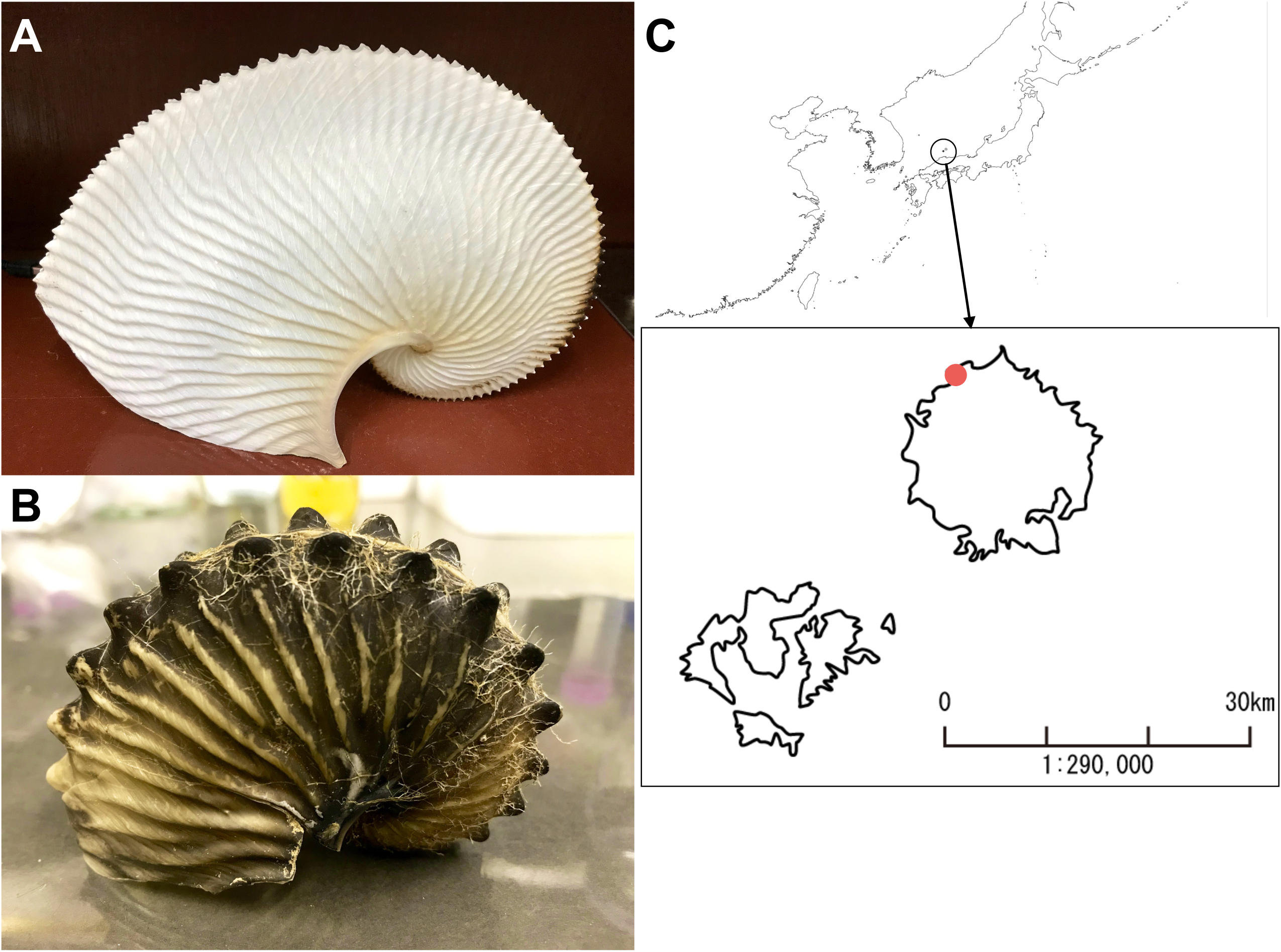
The Argonaut octopuses. A. The shell-like eggcase of *Argonauta argo*. B. The shell-like eggcase of *A. hians*. C. Collect location.

The discrepancy between GenomeScope estimation (1.1Gb, Figure S1) and our actual assembly size (1.34Gb; 1.25Gb non-gap regions) might be caused by the presence of bacterial contamination and/or heterogeneities caused by large insertion and deletion between haploid genomes. However, we only found a very minute amount of bacterial genome contamination in our assembly, indicating that the latter was most likely the main cause of the discrepancy. To assess the completeness of the gene space of the assembly, an analysis using BUSCO v3.0.2 (genome mode) (Simão et al. 2015) was performed by using the provided metazoan data set (metazoa_odb9, n=978), resulting in the recovery of 91.1% of the predicted gene sequences (Table S2). Krait analysis showed that the microsatellite regions account for 4.6% of the genome, with dimer and trimer regions accounting for more than 85% (Figure S2, Table S3).

The high-quality and relatively high level of completeness of our genome assembly, as shown by the statistics we presented above will allow us to address some lingering questions on Cephalopod biology and evolution at the genetic and genomic levels. For example, future studies might utilize the microsatellite regions, which compose a part of the repeat regions in the genome, as individual markers because of the large polymorphisms within individuals, or as markers for paternity analysis of egg-masses.

### Ancient gene clusters in the cephalopods: HOX, Parahox, and Wnt genes

The improved contiguity of our genome assembly confirmed the presence of a Hox cluster. A large Hox cluster of nine Hox genes on four separate scaffolds was recovered in the *A. argo* genome, totaling to a length of at least 18 Mb (Figure 2). Three of the nine Hox genes are not presumed to be gene models, but we have used the Homeobox domain sequence to confirm that they are indeed present on the scaffold and that they are indeed hox genes compared to other Lophotrochozoa genes (Figure S3, Figure S4). *Hox2*/*proboscipedia* (*pb*) was not found, as in squid genomes (Belcaid et al., 2019; da Fonseca et al. 2020) except *Nautilus* (Zhang et al. 2021). We also could not find *Hox4*/*Deformed* (*Dfd*), which is similar to *O. bimaculoides* (Albertin et al., 2015), and thus probably a common feature in benthic octopods (Figure S3). There are at least 10 ORFs inserted among several different Hox genes (3 between *Scr* and *Antp*, 7 between *Lox4* and *Post2*). Interestingly, no homolog was found in other organisms, including even the giant squid *A. dux* (da Fonseca et al., 2020), for any of these ORFs (Table S4).

**Figure 2.**
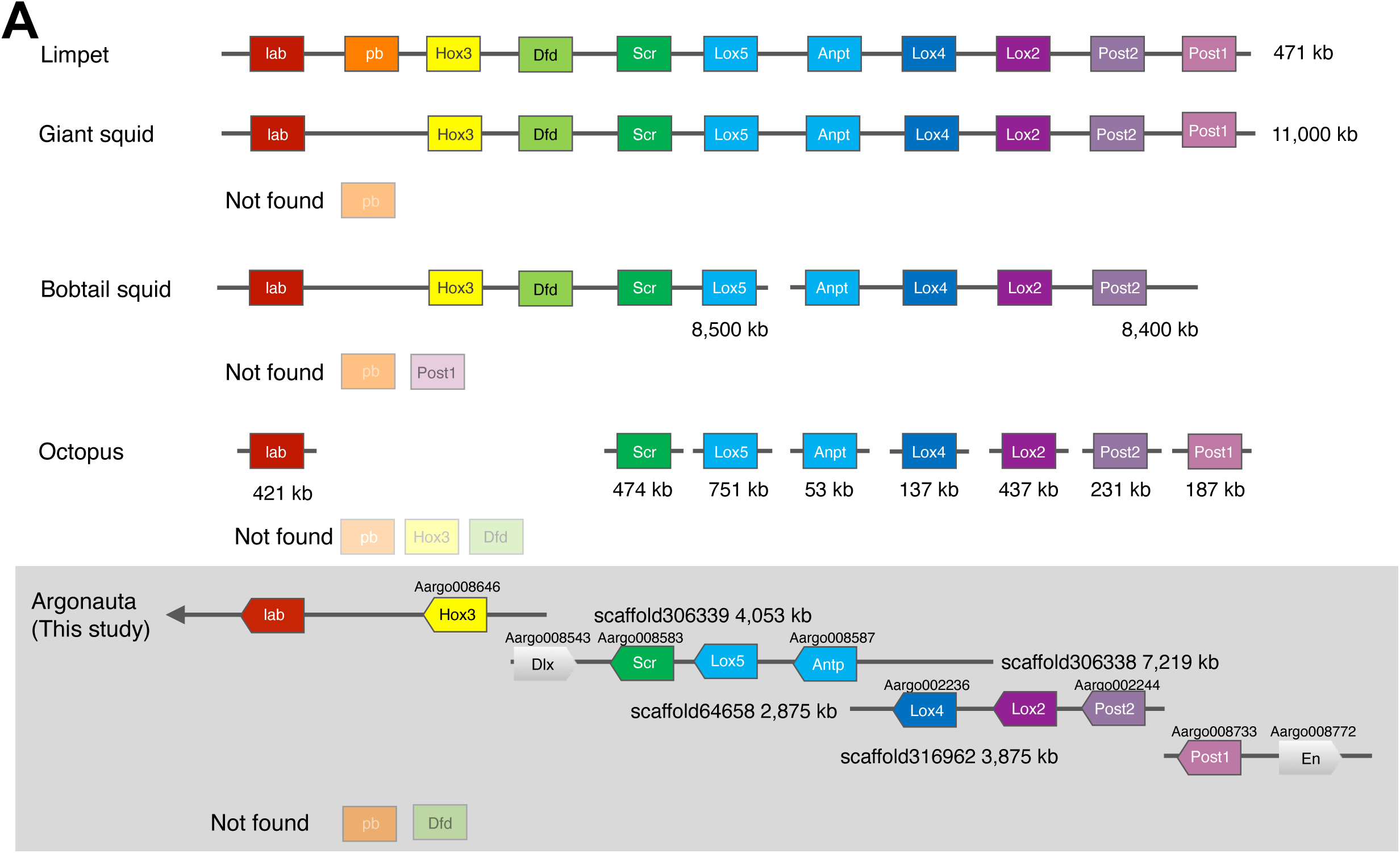
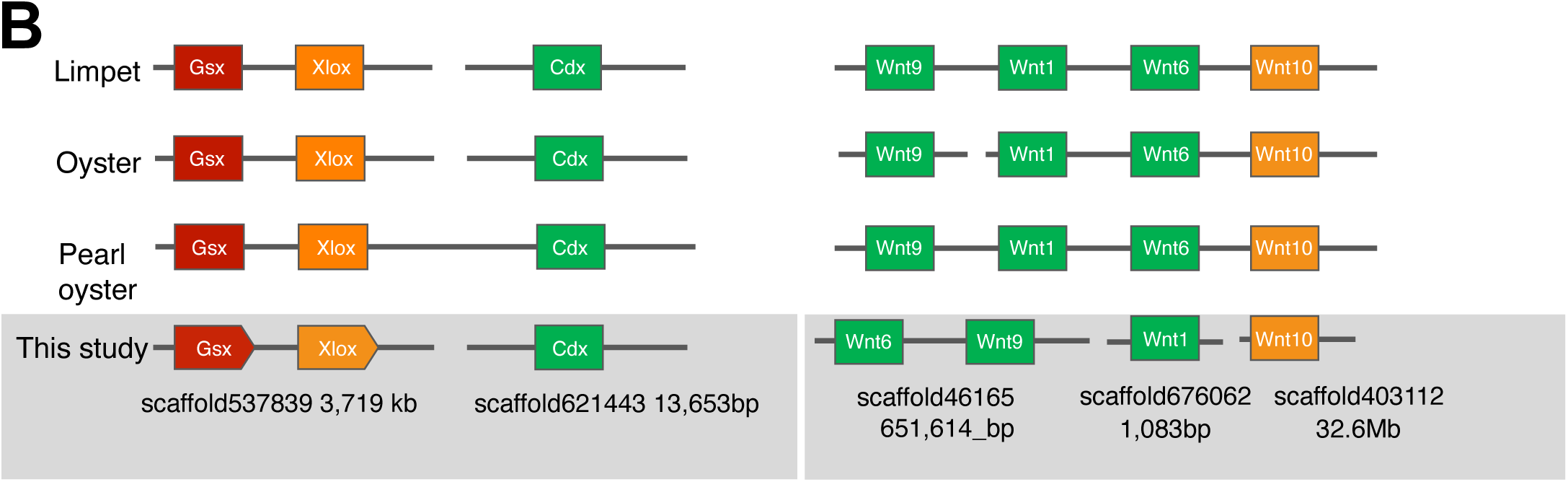
Schematic representations of Hox/Parahox/Wnt clusters. A. Simplified classification of the Hox cluster genomic organization of the cephalopods with the genome sequenced. Scaffold number and length are shown for the *A. argo* genome. The gene model IDs of each gene are shown above each box. The sequences of the homeobox region were confirmed from scaffold for those gene IDs not listed. Hox2/pb and Hox4/Dfd were also not found in the *A. argo* genome as in the *O. bimaculoides* genome. B. Simplified classification of the Parahox and Wnt cluster genomic organizations of molluscs with the genome sequenced. Scaffold number and length are shown for the *A. argo* genome.

Hox clusters are usually found in contigs of about 100 kb in vertebrates and >1,000 kb in invertebrates (Powers et al., 2000; Wagner et al., 2003). Meanwhile, the octopus Hox gene cluster is apparently fragmented, and the genes are present separately on its genome one by one (Albertin et al., 2015), unlike most other bilaterian genomes (Duboule, 2007). Intuitively, this finding seems to be in accordance with the staggered, non-colinear expression pattern of Hox genes in Cephalopods (Lee et al., 2003; Wollesen et al., 2018). However, our finding of a Hox cluster in the genome of *A. argo*, albeit incomplete, suggests that fragmentation of the cluster is probably a feature limited to benthic octopods (or even a possible artifact of the genome assembly process of *O. bimaculoides*).

The presence of ORFs located among several Hox genes in the genome of *A. argo* is also intriguing, since it might indicate that the Hox cluster is actually breaking at the place where the intervening genes are located. The Patellogastropod limpet *Lottia gigantea*, another member of the shelled mollusks (Conchifera), was found to have a typical invertebrate Hox cluster spanning 471 kb with no intervening ORFs among any of its Hox genes (Simakov et al. 2012). Meanwhile, recent findings indicate that although the genome of the basal Cephalopod *N. pompilius* contains a complete set of molluscan Hox genes, they are not located together in a cluster, but are divided in 7 contigs (Zhang et al. 2021). On the other hand, the Hox genes in another Cephalopod, the giant squid *A. dux*, are apparently arranged into a disorganized cluster with insertions of intervening non-Hox genes among cluster members (da Fonseca et al., 2020). However, we found no apparent homology or synteny between any of the intervening ORFs of *A. argo* and those of *A. dux* (Table S4). The acquisition of putative ORFs inside the Hox cluster of *A. argo* is probably an indication of a situation not dissimilar to what was proposed for the fruitfly *Drosophila melanogaster*, which Hox cluster is split into two complexes, with the presence of non-homeotic genes in between (Von Allmen et al., 1996; Wagner et al., 2003; Robertson and Mahaffey, 2017), although *Drosophila* still maintained its colinear expression pattern (Graham et al., 1989; Gaunt, 2015). However, the Hox cluster break in *Drosophila* is most likely a lineage-specific feature, since another model insect, the genome of the beetle *Tribolium castaneum* are intact (Von Allmen et al., 1996; Tribolium Genome Sequencing Consortium, 2008; Shippy et al., 2008). When considered altogether, it seems that the splits and breaking offs of Hox cluster could be a symplesiomorphic feature of the Cephalopod genome, but with the actual “Hox de-clustering” processes happened lineage-specifically. This might explain why Cephalopods do not exhibit typical invertebrate Hox cluster arrangement seen in, for example, the limpet *L. gigantea*.

The Extended Hox complex (*Hox* genes plus *Evx, Mox*, and possibly *Dlx*) is also a common feature in bilaterian genomes (Montavon, 2015). In vertebrate genomes, the complex is shown to be linked to the EHGbox (*En, Hb9*, and *Gbx*) and NKL gene groups (*Msx, Emx*, etc.) and form a supercluster (Garcia-Fernàndez, 2005). In the *A. argo* genome, we found, probably for the first time in Spiralia, a linkage among *Dlx, Engrailed* (*En*), and the Hox genes (Figure 2). In the genome of the giant squid *A. dux, Dlx* and *En* were found in different scaffolds with no linkage to the Hox cluster whatsoever (Table S4). However in *A. argo, Dlx* was found to be located anterior to Scr relative to their positions to the Hox genes (Hox cognate group4), while *E*n was found to be located posterior to *Post1* (Figure 2), and thus reversing the presumed ancestral state (Garcia-Fernàndez, 2005). Although the possibility of their reinsertions into the Hox group cannot be ruled out, this may indicate that the presence of the Extended Hox group is probably conserved in modern cephalopods, although the constraint to preserve gene order is probably relatively weak. The weak constraint in preserving gene order could also explain the “Hox de-clustering”, which characteristics include insertions of ORFs in intergenic regions, observed in Cephalopods.

We also found the presence of the ParaHox cluster, an evolutionary sister complex of the Hox cluster, in the genome of *A. argo* (Figure 2). The ParaHox cluster, which consists of the *Gsx, Xlox*, and *Cdx* gene families, are transcription factors involved in the anterior-posterior development during early embryogenesis of bilaterians (Brooke et al, 1998; Garstang and Ferrier, 2013). The ParaHox cluster is usually found intact in the genomes of Deuterostomes except sea urchin and Ascidians (Garstang and Ferrier, 2013)). However, in Lophotrochozoans, such as the annelid *Platynereis dumerilii* and the limpet *L. gigantea*, only *Gsx* and *Xlox* are clustered together, with *Cdx* broken off and thus unlinked in the genome. The ParaHox cluster of *A. argo* were found to conserve the structure of a typical Lophothrocozoan cluster, similar to those reported in *Nautilus* (Huang et al. 2021) and the octopus (Li et al., 2020). Although further study is still needed, the highly conserved nature of the ParaHox clusters among Cephalopoda, mollusks, and even Lophotrochozoans, indicates a possible presence of an evolutionary constraint to conserve the cluster’s presence and arrangement in the genome, after the breakage of *Cdx* from *Gsx* and *Xlox*.

Similarly, an older gene cluster that is widespread in the animal kingdom is Wnt. Most genomes of bilaterians have a common cluster, *wnt9-wnt1-wnt6-wnt10*, or parts of this cluster (Huang et al., 2021). This ancestral cluster of *wnt*s is thought to originate in the evolution of the common ancestor of cnidarians and bilaterians (Janssen et al., 2010; Holstein 2012). In other shelled mollusks (i.e. Conchifera) such as the rock oyster *Crassostrea gigas* and the Japanese pearl oyster *P. fucata*, the limpet *L. gigantea*, and also in *O. bimaculoides*, the *wnt1-wnt6-wnt10* cluster was conserved (Du et al., 2018a), with *L. gigantea* and *P. fucata* seemingly retaining some of the basal lophotrochozoan / protostome *wnt* paralogs (Cho et al., 2010; Setiamarga et al., 2013). In this study, we also confirmed the linkage of *wnt6-wnt9* in *A. argo* (Figure 2), besides the standard Conchiferan cluster. This suggests that *A. argo* probably also derived this arrangement from the basal metazoan form of Wnt gene orientation and clustering (*wnt9-wnt1-wnt6-wnt10*). Meanwhile, we also observed the lack of *wnt3* and *wnt8*, which seems to be lost in the ancestral protostomes / lophotrochozoans and in ancestral Conchiferans, respectively (Janssen et al., 2010; Setiamarga et al., 2013; Liu et al., 2018; Bai et al., 2020; Wang et al., 2021).

### Tandem gene duplications of gene clusters related to pelagic lifestyle

Our *A. argo* genome assembly, which is of sufficiently better quality than those of previous octopods, allowed us to investigate the existence of tandem gene arrangements. Our searches found two gene clusters, the Reflectins and the Tyrosinases (Figure 3, 4). Both are highly expressed in the first arms of the organism (Figure 4).

**Figure 3.**
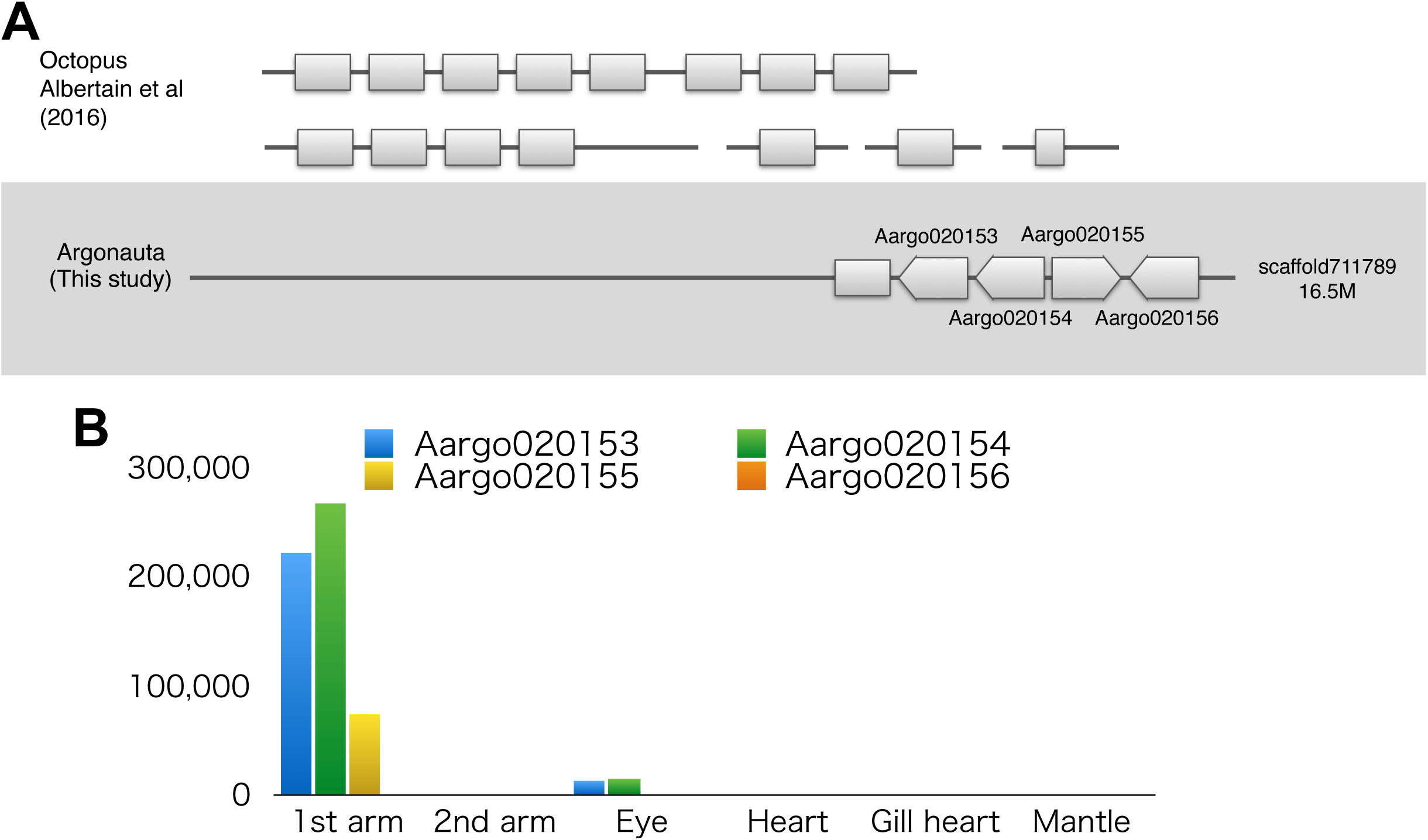
Schematic representations of reflectin clusters. A. Reflectin clusters of the octopuses. B. Gene expression levels of *A. argo* reflectins.

**Figure 4.**
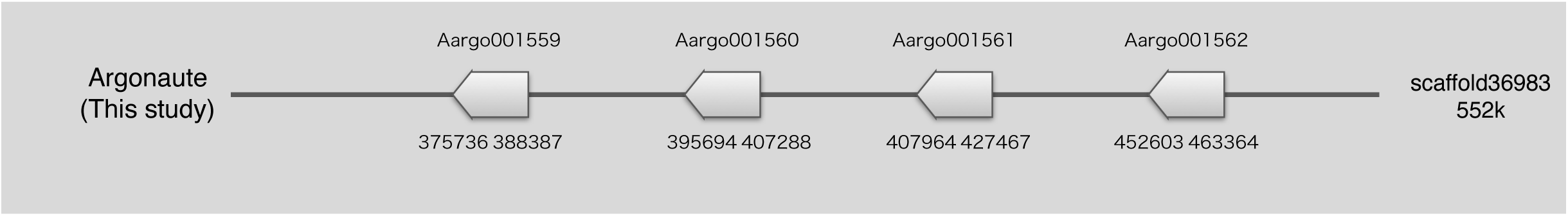
Schematic representations of tyrosinase clusters.

Four tandemly arranged gene models of the octopus *reflectin*/*tbc1* domain family (Aargo020153-6) and one possible ORF recovered by a BLAST search in a single scaffold were found in *A. argo* genome (Figure 3). Phylogenetic analysis showed that the three gene models are monophyletic in *A. argo*, and form a monophyletic clade together with sequences of *E. scolopes*, which also formed a monophyletic clade (Figure S5). The translated sequences of three of the four gene models have at least five of the so-called “Reflectin motifs” (M/FD(X)_5_MD(X)_5_MDX_3/4_) (Levenson et al. 2017, Figure S6). With only 23 nucleotide substitutions, regardless of codon positions, the CDS of the tandemly duplicated *reflectin* genes in *A. argo* match each other sequences at 97%, covering 760 bp (Figure S7). Meanwhile, it is also enticing to suggest that the only gene model with a different sequence, Aargo020154, was inverted to the rest of the genes.

There are two possible causes to explain which duplicated genes are conserved to form gene clusters: either high level expression are favored and thus retaining duplicated genes would help to increase transcript number, or the multiple copies are conserved under different selection pressures as a result of subfunctionalization (Lynch and Force, 2000; Hahn, 2009; Morel et al., 2015; Hallin and Landry, 2019; Song et al., 2020; Ascencio et al., 2021). It has also been pointed out that the duration of concerted evolution can be influenced by selection for a certain dosage of a gene product, as gene conversion leading to highly similar sequence retentions can be advantageous when there is a selection for higher expression level of that particular gene product, or disadvantageous when divergent gene duplicates are advantageous (Sugino and Innan, 2006). Transcriptome analysis shows that in *A. argo, reflectin* is very highly expressed in the 1st arm and eye, and it seems to be transcribed by the three genes (Figure 3B). Therefore, this could be evidence supporting the hypothesis that the cause of gene retention was to have a high level of expression. This concerted evolution may also be the reason why the Cephalopod reflectin formed monophyletic clades with members of the clusters within each species (Figure S5).

Then, what would be the function of the highly expressed *reflectin*? *reflectin* is found only in Cephalopods, and the function of the protein products were shown to be related to camouflage by reflecting and refracting light in the surrounding environment (DeMartini et al., 2015). Expressed proteins fill the lamellae of intracellular Bragg reflectors, allowing individuals to exhibit dynamic iris and structural color changes (Crookes et al. 2004). Several tandemly-arranged *reflectin* gene clusters have been found in the genome of *E. scolopes*, with the dominant *reflectin* transcripts are almost exclusively expressed in the light organ, eyes, and skin, and thus probably consistent with the development of symbiotic fluorescent organs specifically evolved in this lineage (Belcaid et al. 2019). However, although in *E. scolopes*, the symbiotic luminous organs are important for countershading and survival, no such organ have been found in any of the argonauts. As a defence mechanism, pelagic cephalopods blend into their surroundings by camouflaging, which is done either through translucence or cryptic coloration. The first arm membranes of the argonauts are always wrapped around the shell, and reflect light by iridescent chromatophores, causing it to look like a mirror. Meanwhile, the giant squid *A. dux* has seven *reflectin* genes and three *reflectin-like* genes on its genome, all except one are clustered on the same scaffold (da Fonseca *et al*. 2020). This non-luminescent deep-sea species has a mirror-like light-reflecting skin for cryptic coloration. These observations probably indicate that the abundantly expressed Reflectin might help the animals to have light-reflecting mirror-like surfaces, which might then play a role in the ability of these species to blend into their surroundings in the open ocean.

A similar pattern of possible gene conversion was observed in the *tyrosinase* gene cluster. Of the nine *tyrosinase* gene models predicted in *A. argo* genome, eight were of the extracellular or secreted (alpha) type, of which four (Aargo001559-62) were found to be tandemly arranged in a single scaffold (Figure 4, Figure S8). Of the four gene models, excluding unaligned regions, similarities of amino acid sequences of the first two (Aargo001559-60) and the last two (Aargo001561-62) are very high, but only 75% similarities between the two gene pairs. However, the four genes shared an almost exact match in a region on the second half of the gene, at around the 520th - 680th aa (Figure S9). The coding DNA sequence (CDS) match rate for this region is 97% with only ca. 60 substitutions, regardless of codon positions (Figure S10). These two pairs of tyrosinases are orthologous to closely related molluscan taxa including the octopus, and form monophyletic groups (Figure S8). This thus suggests that the four *tyrosinase* copies probably underwent gene conversions in two pairs (between Aargo001559 and 1560, between Aargo001661 and 1662) with some partial recombinations among the four genes. Gene expression analysis using Stringtie shows that the four have a common gene expression profile, with high levels of expression in the arms and mantle. Meanwhile, the phylogenetic tree also indicates that the two pairs of the *tyrosinase* genes are apparently orthologous to those found as shell matrix protein-coding genes in Conchiferan mollusks. This finding, i.e., the genes expressed only in the arms belong to different gene clusters than those of other Tyrosinase-coding genes, might indicate that novel gene paralogs originated from previously existing endogenous *tyrosinase* genes were being duplicated and obtained high expression in the arms, possibly used for the calcified eggcase formation, which helps *A. argo* and other argonaut octopods to attain buoyancy and thus their pelagic lifestyle.

### Opsin duplications and change in absorption wavelength related to pelagic lifestyle

Changes in number and sequences of Opsin are thought to be involved in adaptation to visually-guided behavior. We found that *A. argo* possesses five visual pigment genes in its genome: two noncanonical *r-opsins*, one canonical *r-opsins*, two *xenopsin*, and one *rgr/peropsins/retinochromes* (Figure S11-13) (Ramirez et al., 2016). In previous studies on the pygmy squid *Idiosepius paradoxus*, two *r-opsins*, one *xenopsin*, and two *retinochromes* were identified (Yoshida et al. 2015; Ramirez et al. 2016). We also checked if the expression of the opsins are tissue specific, in order to see whether there is any functional differentiation among the duplicated opsins. However, at present, we were unable to confirm such specificity, at least in the organs we examined because the data is present, such as the eye and skin. In fact, almost no study has been conducted on the functions of *xenopsins* and non-canonical *r-opsins* in the Cephalopods. In the future, a thorough gene expression analysis of these genes in different tissues should probably be conducted to resolve this issue.

The presence of two xenopsins in the genome of *A. argo* was apparently not an artifact or assembly error, which means that *A. argo* has an extra copy of *xenopsin* than *I. paradoxus*. The gene models for *xenopsin* in *A. argo* (Aargo004635 and Aargo004636) exist in tandem in the scaffold, albeit with the amino acid sequence being too short to be considered full-length. If we also assume that the two exons of the neighboring Aargo004633 are shared among the three gene models, we can obtain two putative complete Xenopsin proteins. In other words, it makes sense to think that the two xenopsins were probably splicing variants with alternative promoters and shared two exons, which then duplicated and subfunctionalized (Force et al., 1999; Hahn, 2009). *xenopsin* is found to be widespread but exclusively only in protostomes, co-expressed together with *r-opsin* mostly in their ciliary photoreceptor cells (Passamaneck et al., 2011; Vöcking et al., 2017; Rawlinson et al., 2019). Functional studies on *xenopsin* are lacking and we therefore cannot decisively predict its function in *A. argo*. Phylogenetically, *xenopsin* and *c-opsin* are apparently spread exclusively from each other, with *c-opsin* being found exclusively in the photoreceptor cells of deuterostomes / vertebrates, suggesting that *xenopsin*, similar to its deuterostomian counterpart *c-opsin*. is probably involved mainly in phototactic responses and visual functions in protostomes (Döring et al., 2020), including *A. argo, I. paradoxus*, and probably, other Cephalopods.

The two copies of non-canonical *r-opsins* in the genome of *A. argo* is most likely due to the duplication of heterogeneous regions in the assembly, since the sequences matched perfectly. This suggests that there is only a single non-canonical *r-opsin* in the genome of *A. argo*, which is thus similar to *I. paradoxus*, as mentioned previously (Yoshida et al. 2015). At present, the function of this Opsin homolog is still unknown, although previous studies suggest that it’s probably unrelated to visions, although apparently still related to photoreception (Ramirez and Oakley, 2015; Ramirez et al. 2016; Bonade et al., 2020). We found two amino acid substitutions (T118S and Y178F) in the amino acid sequence of the non-canonical R-Opsin of *A. argo* when compared to bovine rhodopsin. T118S was found in both the benthic *O. bimaculoides* and *A. argo*, while Y178F was found only in the latter. Prediction of light absorption wavelength of the non-canonical R-Opsin of *A. argo* indicates that photoreceptions in *A. argo* are probably adapted more to red light than that of the benthic octopus. The extra amino acid substitutions are thus consistent with the ecology of *A. argo*, which lived closer to the sea surface than other cephalopods, indicating that the red-shift may be an adaptation to shallow water light environment.

### The evolution of shell and eggcase matrix proteins through independent recruitments, losses, and domain changes allows *A. argo* to obtain its eggcase and thus its pelagic lifestyle

In this study, we found all of the eggcase matrix protein-coding genes in the genome of *A. argo* (Table S5) as identified by our previous multi-omics study to survey and identify major proteins of the eggcase matrices of two congeneric argonaut octopods, *A. argo* and *A. hians* (Setiamarga et al., 2021b). Exactly congruent to our previous result, most of the proteins are apparently not shared with the shell matrix proteins of Conchiferans, including those of the basal Cephalopoda *Nautilus* (Setiamarga et al., 2021a; Huang et al., 2021), although the genes / proteins themselves are present in the genomes of the Conchiferan mollusks such as the limpet *L. gigantea* (Simakov et al., 2013), the true oyster *C. gigas* (Peñaloza et al., 2021), and the Japanese pearl oyster *P. fucata* (Takeuchi et al., 2016). Meanwhile, the Conchiferan shell matrix protein-coding genes were also mostly found in the genomes of *A. argo* and the shell-less benthic octopod *O. bimaculoides* (Albertin et al., 2015), indicating their retention despite shell loss in the octopod lineage. Interestingly, the genes for eggcase matrix proteins were also found in the genome of *O. bimaculoides*, and thus, when considered altogether, supported our hypothesis suggested previously, saying the argonaut octopods recruited many proteins unrelated to the shell formation and used them for their eggcase (Setiamarga et al., 2021b). However, very interestingly, some proteins related to calcification such as the Pif-like LamG3, seemed to be used at least by *A. hians* (Setiamarga et al., 2021b).

In that previous study, we arbitrarily categorized the Pif-like proteins mostly identified as Conchiferan shell matrix proteins into three paralogous groups, based on three monophyletic clades recovered in the phylogeny (see Figure 5 in Setiamarga et al., 2021b), which were also recovered in this study (Figure 5). We arbitrarily named them Blue Mussel Shell Protein (BMSP), Laminin G3 (LamG3), and Pif, and called them altogether the BMSP/LamG3/Pif proteins. These proteins could be distinguished by their domain combinations. BMSP was first identified as SMPs in the blue mussel *Mytilus galloprovincialis* and *L. gigantea*, respectively (Suzuki et al. 2011; Marie et al. 2017). The protein has one Chitin-Binding (ChtBd) and multiple (three or four) von Willebrand factor type A (VWA) domains. BMSP is present throughout the nacreous layer with dense localization in the myostracum, suggesting its possible role in Conchiferan nacreous layer formation (Suzuki et al. 2011). Meanwhile, Pif proteins, which was originally found in the nacre of *P. fucata*, usually have two types of domains, one VWA and two ChtBd domains, but with a different domain compositions and arrangements (Figure 5) (Suzuki et al. 2009; Setiamarga et al., 2021a, b). *In vitro* functional analysis has shown that it is involved in calcium crystallization (Suzuki et al. 2013).

**Figure 5.**
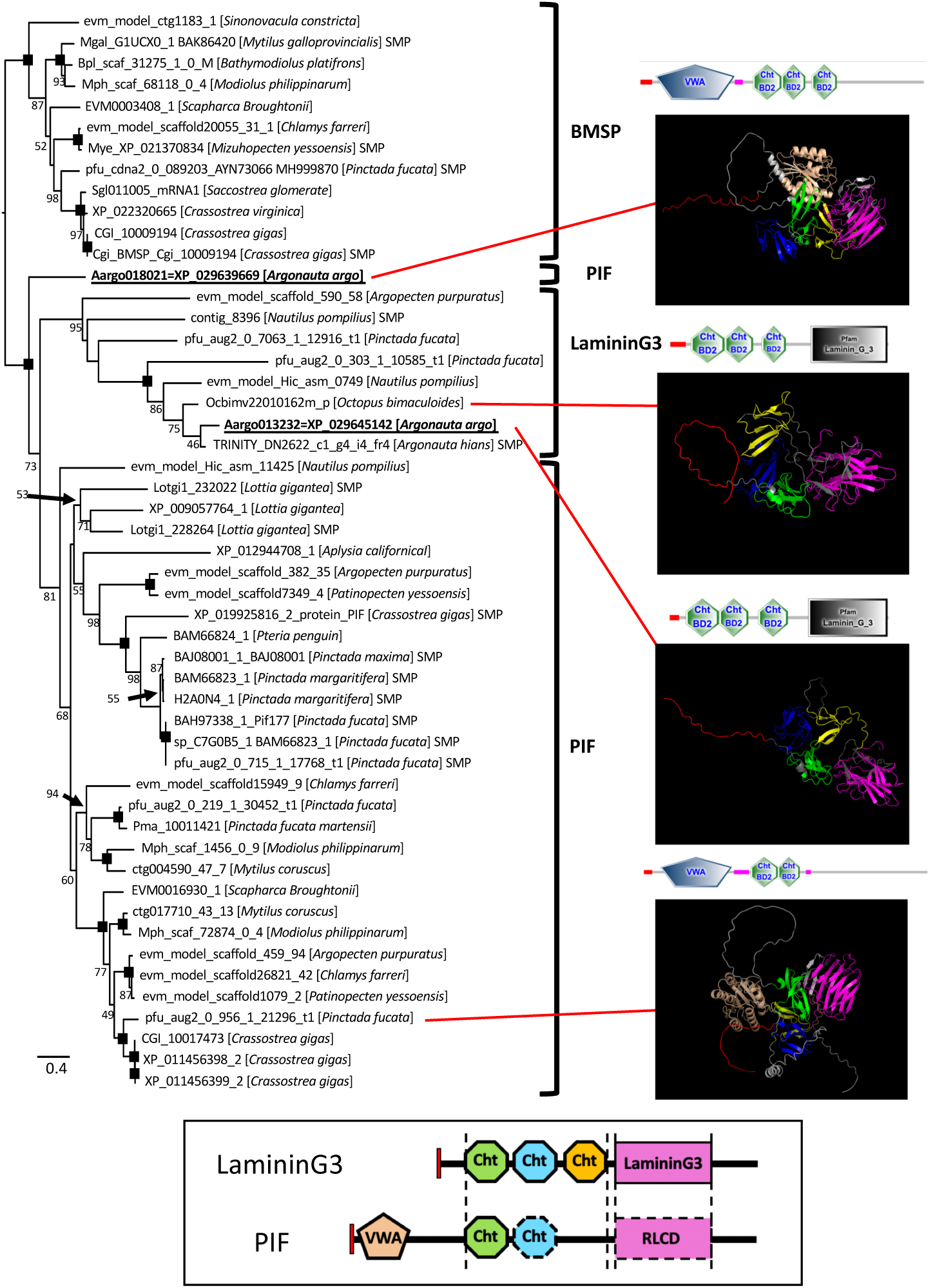
Phylogenetic relationships of Pif/Pif-like/BMSPs of Molluscs and representative 3D protein models. The maximum likelihood tree was estimated under the best fit models (WAG + Γ). Numbers on the nodes are Bootstrap Support (BS) values. BS lower than 41% are shown as “--”, while 100% support is not written. Representative structures of the proteins of the sequences included in the analyses, shown as SMART protein domains, are shown below the trees. Four 3D structural models (PIF; Aargo018021 [*Argonauta argo*], pfu_aug2_0_956_1_21296_t1 [*Pinctada fucata*], LamG3; Aargo013232 [*Argonauta argo*], Ocbimv22010162m_p [*Octopus bimaculoides*]) were estimated with AlphaFold2. Schematic representation of domain structure and 3D structural model were colored each domain characteristic: Signal peptide, red; VWA, pale orange; 1st ChtBd, green; 2nd ChtBd, blue; 3rd ChtBd, yellow; LamG and RLCD, pink.

We did not detect any of the homologs of Pif and BMSP in the eggcase matrix of the argonauts in our previous multi-omics eggcase matrix protein study (Setiamarga et al., 2021b), nor in the shell matrix protein study of *Nautilus* (Setiamarga et al., 2021a). LamG3 was first identified by Marie et al. (2017) as one of the two Pif-like isoforms composed of the one VWA, three ChtBd, and one LamG domains. In both cephalopods, we instead found the last type of Pif homologs (*sensu* Setiamarga et al., 2021b), the LamG3 protein, in both the eggcase matrix of *A. hians* (but not in the eggcase matrix proteome and transcriptome of *A. argo*), and the shell matrix of *Nautilus*. However, very interestingly, differing with the results of our previous multi-omics study (Setiamarga et al., 2021b), in this study, we found the presence of *lamG3* in the genome of *A. argo* (Aargo013232) (Figure 5). Further studies must thus be conducted to assess if the absence of any transcript/protein product of *lamG3* in *A. argo*, despite its presence on the genome, is an artifact caused by the possible non-exhaustiveness of our previous multi-omics study, or if it is not used in the eggcase matrix of *A. argo*, making the eggcases of the two congeneric species different in nature.

At present, however, we are working under the hypothesis that this protein is a key protein for the formation of calcified eggcases in argonaut octopods, because it is one of the putative paralogs of the BMSP/LamG3/Pif-like proteins, which members have been identified as major component of the shell of Conchiferan mollusks. Intriguingly, however, although in the previous multi-omics study we did not find any sequence of *pif* or *BMSP* both in the transcriptome data of all tissues studied and the proteome data of *A. argo*, we found the presence of an intact coding sequence of *pif* in the genome of the species (Aargo018021). More interestingly, an intact *pif* sequence was also found in the genome of *N. pompilius* (Huang et al. 2021), although the sequence was not found in the transcriptome and proteome data of our recent shell matrix proteins multi-omics study of the species (Setiamarga et al., 2021a). The exon-intron structures of each cluster are different and are located at different positions in the genome of both *A. argo* (Figure 5b) and *Nautilus*. Meanwhile, LamG3 has also been shown to be associated with the biomineralization of shells in the pond snail *Lymnaea stagnalis*, although no BMSP nor Pif were apparently found in its shell matrix proteome and transcriptome, although additional studies involving genome analysis of the species is still needed to confirm this observation (Ishikawa et al. 2020). These results seem to thus indicate that the two Pif homologs (*pif* and *lamG3*) were probably already present separately at least in the basal Conchiferan mollusks. However, ancestral Cephalopods even more basal than the *Nautilus* probably lost *bmsp*, retained *lamG3* and *pif*, but use only *lamG3* for the formation of biomineralized shells. Although further confirmation is still needed, *lamG3* was probably recruited independently as an SMP, independently in each lineage leading to terrestrial gastropods and cephalopods. The presence of *pif* in the genomes of *Nautilus* and *A. argo*, and *lamG3* in the genome of *O. bimaculoides*, even though they are not involved in the formation of shells or shell-like structures, is probably because they acquired new functions unrelated to shell formation.

The lack of a typical LamG3 domain in BMSP and Pif have been reported (Suzuki et al., 2013), and domain searches using various tools such as SMART (http://smart.embl-heidelberg.de/, accessed in June 2021), InterProScan (https://www.ebi.ac.uk/interpro/search/sequence/, accessed in June 2021), and Pfam (http://pfam.xfam.org/, accessed in June 2021) seemed to support this notion. In both BMSP and Pif, no domain was detected. Some searches would detect only the repetitive low-complexity domains (RLCD) which designates a possibility that the particular region used to have a domain, but has probably degraded down and thus only recognizable partially at the sequence level (Suzuki et al., 2017; Setiamarga et al., 2021a; b). We also predicted the stereostructures of some representatives of BMSP (*Mytilus galloprovincialis*), Pif (*A. argo, N. pompilius, P. fucata, L. gigantea*), and LamG3 (*A. argo, N. pompilius, O. bimaculoides, P. fucata, L. gigantea, C. gigas*) using Alphafold2 (Jumper et al., 2021), and compared their 3D structures (Figure 5). AlphaFold2 predicts accurate protein 3D structural models. Structural comparisons using such models would allow us to obtain surprising information about the function and evolution of the proteins, unattainable only through sequence comparison and comparative genomics. The models of BMSP/LamG3/Pif proteins predicted by AlphaFold2 indicate that the region where no domain was detected in the proteins (except for the LamG3 proteins) actually still retains enough of its LamG3 domain characteristics (Figure 5). LamG3 domain is a receptor for various extracellular matrix proteins, which function is mediated by the calcium ion (Tryggvason 1993; Yurchenco et al. 1993; Yu and Talts 2003; Klees et al., 2008; Suzuki et al., 2017). This thus might explain the usefulness of the domain not only for the formation of calcified structures but also for other functions, while at the same time also suggests the unnecessity of the organisms compared to retain or use all of the BMSP/LamG3/Pif protein homologs for the same function, which might thus also explain why Cephalopods (*Nautilus*) only use LamG3 for their shell formation, and why the argonauts also re-recruited this protein to form their eggcase.

## Conclusion

Until very recently, studies on the evolution of Cephalopoda lacked insights from genomic perspectives. However, recent genome data publications of various species have remedied this. In this study, we present a genome assembly of *Argonauta argo*, which provides significant insight into the genetic and evolutionary background of the adaptation to the pelagic environment, such as the evolution of the visual proteins Opsin and Reflectin, and the shell matrix protein Tyrosinase. The improved quality of the genome assembly also allowed us to identify the presence of sexually highly polymorphic regions, which would be useful in future studies aiming at the elucidation of the genetic underpinnings of extreme male-female dimorphisms in the species. The pronounced sexual dimorphism probably evolved as an adaptation to holopelagic life in the open ocean with few male-female encounters. Besides that, the improved contiguity of the genome assembly confirmed the presence of several gene clusters including both deeply conserved ones, such as Hox, ParaHox, and Wnt, and unique ones that might be involved in evolutionary novelty.

The newly obtained draft genome sequence also allowed us to hypothesize about the evolution of some major shell matrix proteins related to calcification, seemingly re-recruited in the formation of the eggcase, which was impossible to do in our previous multi-omics-based studies. We also were able to corroborate our previous report based on a multi-omics study on the eggcase matrix proteins. In this study, we found all of the eggcase matrix proteins previously identified, while at the same time, also found the presence of LamG3 in the genome of *A. argo* (and *O. bimaculoides*), which was found as one of the eggcase matrix proteins of *A. hians* but not in *A. argo* in our previous multi-omics study. We also found an ortholog of the Pif coding gene in the genome of *A. argo*, besides in the recently published genome of *N. pompilius*. Combined with the protein structure prediction using Alphafold2, we thus were able to build a hypothesis about how BMSP/LamG3/Pif proteins evolved. In our hypothesis, the BMSP/LamG3/Pif proteins are key proteins for the formation of calcified external structures, including the eggcase. Therefore, the presence of *pif* in the genomes of *Nautilus* and *A. argo* and *lamG3* in the genome of *O. bimaculoides* might explain the usefulness of LamG3 domain for the formation of calcified structures, which might thus explain why the argonauts also re-recruited LamG3 protein, although not necessarily Pif and BMSP, to form their eggcases.

## Materials and Methods

### Sampling, Sequencing, and Genome Size Estimation

The *A. argo* DNA used for sequencing was derived from a single female provided by bycatch caught in the fixed nets set along the coast in Oki Island Town, Shimane Prefecture, Japan (36°17’20.6”N 133°12’46.4”E). Pieces of the gonad (ovary) were collected from an individual female specimen collected in 2018. The shell is registered as a collection of The University Museum, The University of Tokyo in Tokyo, Japan (Voucher No. RM33391). Genomic DNA was extracted from the ovary using the QIAGEN Genomic-tip kit. Pooled DNA was used for the preparation of three paired-end and three mate-pair (3, 6, 10, and 15 kbp insert size) libraries, that were sequenced on an Illumina HiSeq 2500 at the National Institute of Genetics, Japan with supports by Platform for Advanced Genome Science (PAGS) (Table S6, S7).

Pieces of the mantle, arm membrane of the first arm, and 2nd arm tip were obtained from the same single individual to genomic DNA. Eyes, hearts, and gill hearts were sampled from different individuals of *A. argo*. Six transcriptomes of *A. argo* were obtained and raw data statistics are provided in Table S6. Total RNA was extracted from the tissue samples using Trizol (Invitrogen) followed by an on-column DNaseI treatment using the RNeasy mini kit (Qiagen). The RNA acquoliot was stored at –80ºC until further transcriptome analyses. The *A. argo* genome size and heterozygosity were assessed with GenomeScope v2.0 (Ranallo-Benavidez et al. 2020), based on the quality-filtered Illumina reads. A heterozygosity rate of 1.44% was estimated from the 32-mer-based assessment of the *A. argo* genome (Supplementary Figure S1). Complete microsatellite sequences were estimated and visualized with Krait v1.3.3 (Du et al. 2018b).

Raw read sequence data will be available in the DNA Data Bank of Japan (DDBJ). We are willing to share our raw data before the publication of the original paper on the assumption that it will be done as a collaborative research.

### De Novo Genome Assembly and Annotation

Using the predicted 1.1 Gb genome size estimate of the *A. argo*, the total raw sequence coverage of Illumina reads was 201× (pair-end reads, 3 kb, 6 kb, 10kb, and 15 kb mate-paired libraries). To reconstruct the mitogenome, we performed contig assembly (-n 200) with Platanus v1.2.4 (Kajitani et al. 2014) using the paired-end data. Contigs annotated as mitochondrial sequences were extracted by using the mitogenome data of a closely related species, *A. hians* (NC_036354), as the query for BLASTn homology search. After assembling the contigs, both ends of the resulting single contig were manually confirmed to overlap, and redundant parts were removed to complete the full circular mitogenome.

The pair-end sequence reads (PE600) after adapter trimming were assembled using De Bruijn graph assembler, Platanus-allee v. 2.2.2 (Kajitani et al. 2019). The basic algorithm of the Platanus-allee v2.2.2 is based on the arrangement of two independently assembled sequences derived from each haplotype of the corresponding two homologous chromosomes. Contig assembly was performed using only the PE library, and then scaffolding and gap closure were performed using all libraries. Assembly statistics by Platanus v222 was shown in Table S8.

Gene prediction models were generated using custom-made annotation pipeline as in (Inoue et al. 2021). In brief, this pipeline combines RNA-seq-based prediction results, homology-based prediction results for related species, and ab initio prediction results using in-house dynamic program. RNA-seq based prediction utilized both the assembly-first method and the mapping-first method. For the assembly-first method, RNA-seq data were assembled using Trinity (Grabherr et al. 2011) and Oases (Schulz et al. 2012). Then, assembled contigs were splice-mapped with GMAP (Wu et al. 2005). For the mapping-first method, RNA-seq data were mapped to genome scaffolds and genes were predicted using HISAT2 (Kim et al. 2019) and StringTie (Petea et al. 2016). In terms of homology-based prediction, amino acid sequences of *Octopus vulgaris* (Zarrella *et al*. 2019), *Octopus bimaculoides* (Albertin *et al*. 2015), *Architeuthis dux* (da Fonseca *et al*. 2020), *Crassostrea giga* (Zhang *et al*. 2012), and *Mizuhopecten yessoensis* (Wang *et al*. 2017), were spliced-mapped to genome scaffolds using Spaln62, and gene sets were predicted. For ab initio prediction, raining sets were selected from RNA-seq based prediction results and AUGUSTUS (Stanke et al. 2003) and SNAP (Korf et al. 2004) were trained and used for prediction. Predicted results of each tool are shown in Table S9 and as a final result, 20,293 protein coding genes were predicted (Table S9). Predicted genes were evaluated using BUSCO v3.0.2 (protein mode) (Simão et al. 2015) and resulted in 97.0% complete gene marked, suggesting high accuracy of the annotation (Table S10). This goes beyond the cephalopod genomes sequenced so far, and is comparable to high quality mollusc genomes (Table S11).

### Phylogenetic Analysis

Phylogenetic analyses were conducted on a total of five gene families obtained in this study (*Hox, reflectin, tyrosinase, opsin, bmsp/lamg3/pif proteins*). To build single-gene trees based on orthologs, we performed webBLAST search using *A. argo* protein sequences translated from the gene sequences. Sequences for the phylogenetic tree were collected from Genbank to cover the whole Lophotrochozoan clade. To perform multiple alignments of protein sequences, we utilized the online version of MAFFT v7.487 (Katoh et al. 2002; https://mafft.cbrc.jp/alignment/software/; accessed in August 2021), followed by the removal of ambiguously aligned sites using the online version of trimAl_v1.4beta (automated option) (Capella-Gutiérrez et al. 2009; http://phylemon2.bioinfo.cipf.es/index.html; accessed in August 2021). Maximum likelihood phylogenetic inferences were executed on the software RAxMLGUI v2.0.5 (Silvestro et al. 2012; Stamatakis 2006) the rapid tree search setting with 1000 bootstrap replications under the best fit models (BMSP/LamG3/Pif proteins = WAG + Γ, Hox = LG + Γ + I, Reflectin = JTT + Γ + F, Tyrosinase = LG + Γ + I). The best fit models were inferred using MEGA X (Kumar et al. 2018). Obtained trees were visualized with FigTree v1.4.2 (Rambaut 2009).

For Opsin, sequences from other metazoans were collected from GenBank and Ensembl databases. Multiple sequence alignments of protein sequences were also performed by MAFFT. The best fit models were inferred using Modeltest (Darriba et al. 2020). Maximum likelihood phylogenetic inferences were executed on the software IQ-TREE (ref) the tree search setting with 1000 bootstrap replications under the best fit models (LG+G4: Best-fit model according to Bayesian Information Criterion (BIC) for c-opsins, LG+F+I+G4: BIC for r-opsins). The trees were visualized with FigTree v1.4.2 (Rambaut 2009).

## Acknowledgments

Samples were obtained with the courtesy of Mr. Minoru Yoshida and his colleagues of Yoshida Suisan, Oki Islands. Drs. Noriyoshi Sato of Tokai University and Hiroki Ono of Shimane University contributed to sample collections. Computations were partially performed on the NIG supercomputer at ROIS National Institute of Genetics, Shizuoka, Japan. We are grateful to Drs. Kazutoshi Yoshitake and Takeshi Kawashima for their help on how to use the Singularity module on the NIG supercomputer system. DHES, HK, and TS would like to thank members of Setiamarga Lab at National Institute of Technology, Wakayama College, and Sasaki Lab at The University Museum of The University of Tokyo, for their constant support during the commencement of the study.

The work was supported by Human Frontier Science Program grant (RGP0060/2017) to MAY, Japan Society for the Promotion of Science (JSPS) KAKENHI Grant Number 18K06363 (Grant-in-Aid for Scientific Research (C))] to MAY and DHES and 16H06279 (PAGS) to AT and TI. DHES was also supported by the FY2016 Research Grant for Chemistry and Life Sciences (The Asahi Glass Foundation), and partially supported by the FY2017 Research Grant for Zoology (Fujiwara Natural History Research Foundation) and the FY2019 Grant-in-Aid for Challenging (Exploratory) Research (Grant number: 19K21646) (awarded to TS (PI) and DHES (Co-PI)). KI was partially supported by the Platform Project for Supporting Drug Discovery and Life Science Research from the Japanese Agency for Medical Research and Development (AMED). We also thank the faculty of Life and Environmental Science at Shimane University for help in financial support for publishing this report.

## Figure legends and Tables

Figure S1 GenomeScope result.

Figure S2 Microsatellite types found in the Argonaut genome.

Figure S3 Molecular phylogenetic tree of the Hox genes. The maximum likelihood phylogenetic tree inferred under the LG + Γ + I model with 1000 bootstrap replicates. Hox genes of *Argonauta argo* are marked with a black arrow. Abbreviations: Nuctum: *Nucula tumidula*, Cragig: *Crassostrea gigas*, Pecmax: *Pecten maximus*, Gibvar: *Gibbula varia*, Lotgig: *Lottia gigantea*, Apcal: *Aplysia californica*, Eupsco: *Euprymna scolopes*, Octbim: *Octopu bimaculoides*, Naupom: *Nautilus pompilius*, Acacri: *Acanthochitona crinite*, Antent: *Antalis entails*, Glympell: *Gymnomenia pellucida*, Alivir: *Alitta virens*, Linana: *Lingula anatine*: Dromel: *Drosophila melanogaster*, Braflo: *Branchiostoma floridae*: Caeele, *Caenorhabditis elegans*.

Figure S4 Alignment of Hox genes recovered in the scaffolds but not in the gene models.

Figure S5 Reflectin phylogenetic tree

Figure S6 Reflectin alignment with reference to repetitive reflectin motifs

Figure S7 Reflectin alignment to show gene conversion

Figure S8 Tyrosinase phylogenetic tree. The maximum likelihood phylogenetic tree inferred under the LG + Γ + I model with 1000 bootstrap replicates. Numbers on the nodes are Bootstrap Support (BS) values. BS lower than 41% are not shown, while 100% support is shown as a black square. Tyrosinase of *Argonauta argo* are marked with underlined. Three type of tyrosinase are shown secreted (α), cytosolic (β) and membrane-bound (γ) subclasses.

Figure S9 Tyrosinase alignment at amino acid level

Figure S10 Tyrosinase alignment to show gene conversion

Figure S11 Opsin alignment and unique amino acid changes of argonaut based on bovine rhodopsin

Figure S12 Phylogenetic tree of RGR and rhabdomeric opsins

Figure S13 Xenopsin phylogenetic tree

Figure S14-16 Protein structure of Pif/Pif-like/BMSPs. The schematic representation of three proteins of Pif/Pif-like/BMSPs are shown within the box frame. Conserved domains within each protein are predicted in SMART, and the 3D structural models were estimated with AlphaFold2. The domains regions, which distinguished by the domain prediction and its conserved alignment regions, are marked different color: Signal peptide (red), 1st VWA (deep orange), 2nd VWA (ocher), 3rd VWA (pale orange), 4th VWA (orange), 1st ChtBd (green), 2nd ChtBd (blue), 3rd ChtBd (yellow), LamG and RLCD (pink).

Table S1 Assembly comparison among molluscan genomes

Table S2 Assembly comparison based on BUSCO scores (genome mode, metazoa_odb9, n=978)

Table S3 Krait estimation of the microsatellite regions

Table S4 Comparison of ORFs in the Hox cluster between blue mussels and giant squid

Table S5 Proteome list

Table S6 List of *A. argo* genome sequencing data

Table S7 List of *A. argo* RNA-seq sequencing data

Table S8 Assembly statistics by Platanus v222

Table S9 Gene prediction models using custom-made annotation pipeline with transcriptomic data

Table S10 Gene model comparison based on BUSCO scores (protein mode, metazoa_odb9, n=978)

Table S11 Gene model comparison among molluscan genomes

## Notes

### Competing Interest Statement

The authors have declared no competing interest.

