## Supplementary material for "Gene recruitments and dismissals in argonaut octopus genome provide insights to pelagic lifestyle adaptation and shell-like eggcase reacquisition": Figure S

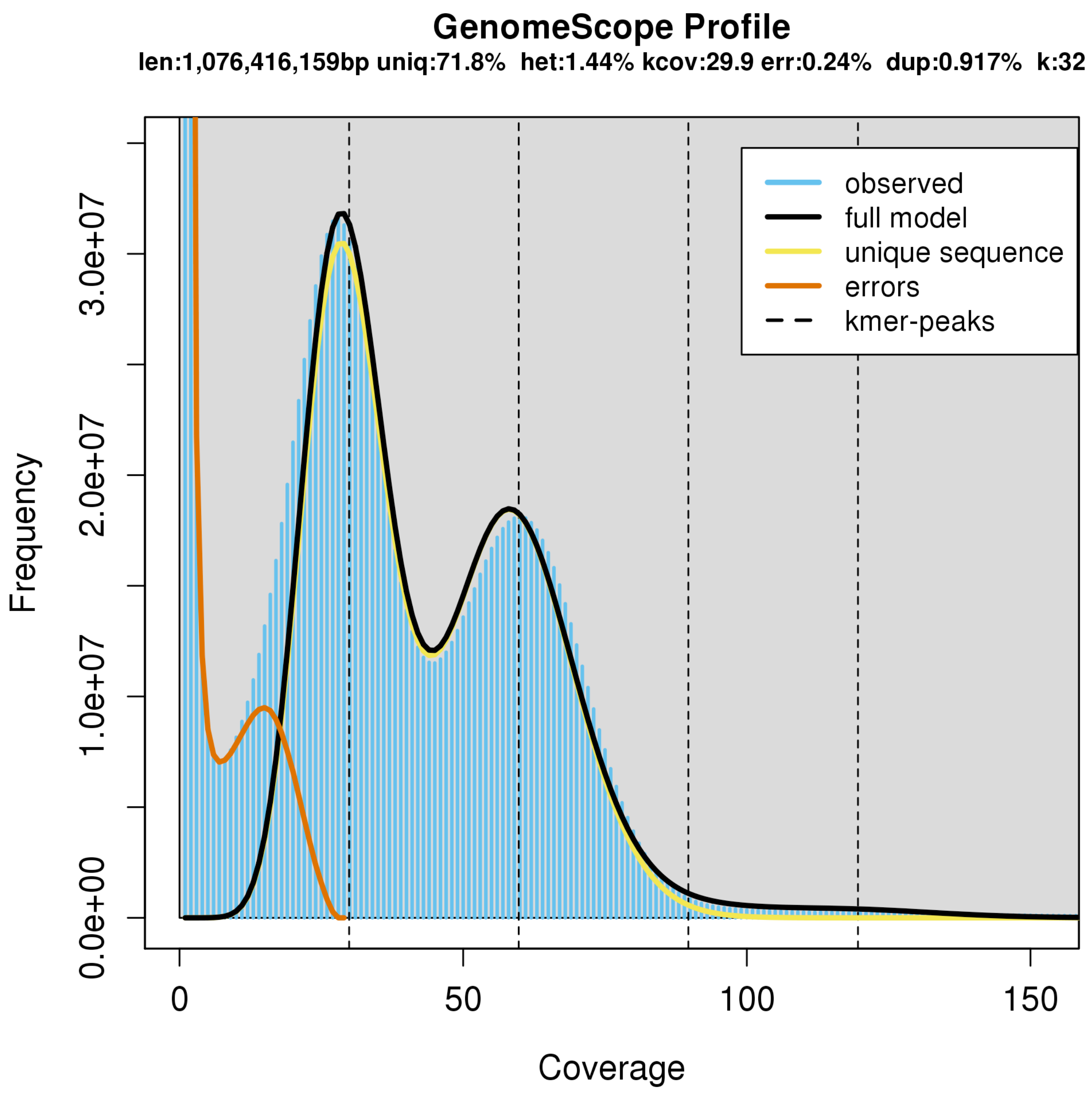


Figure S1 K-mer based genome size estimation

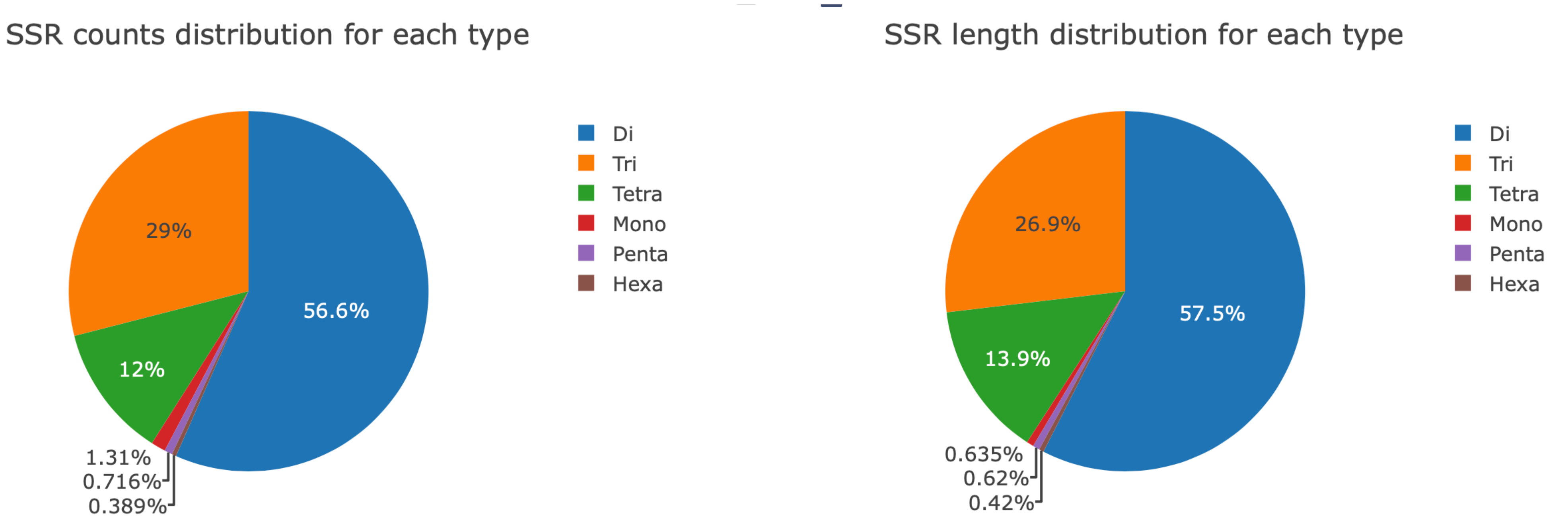


Figure S2 Microsatellite types

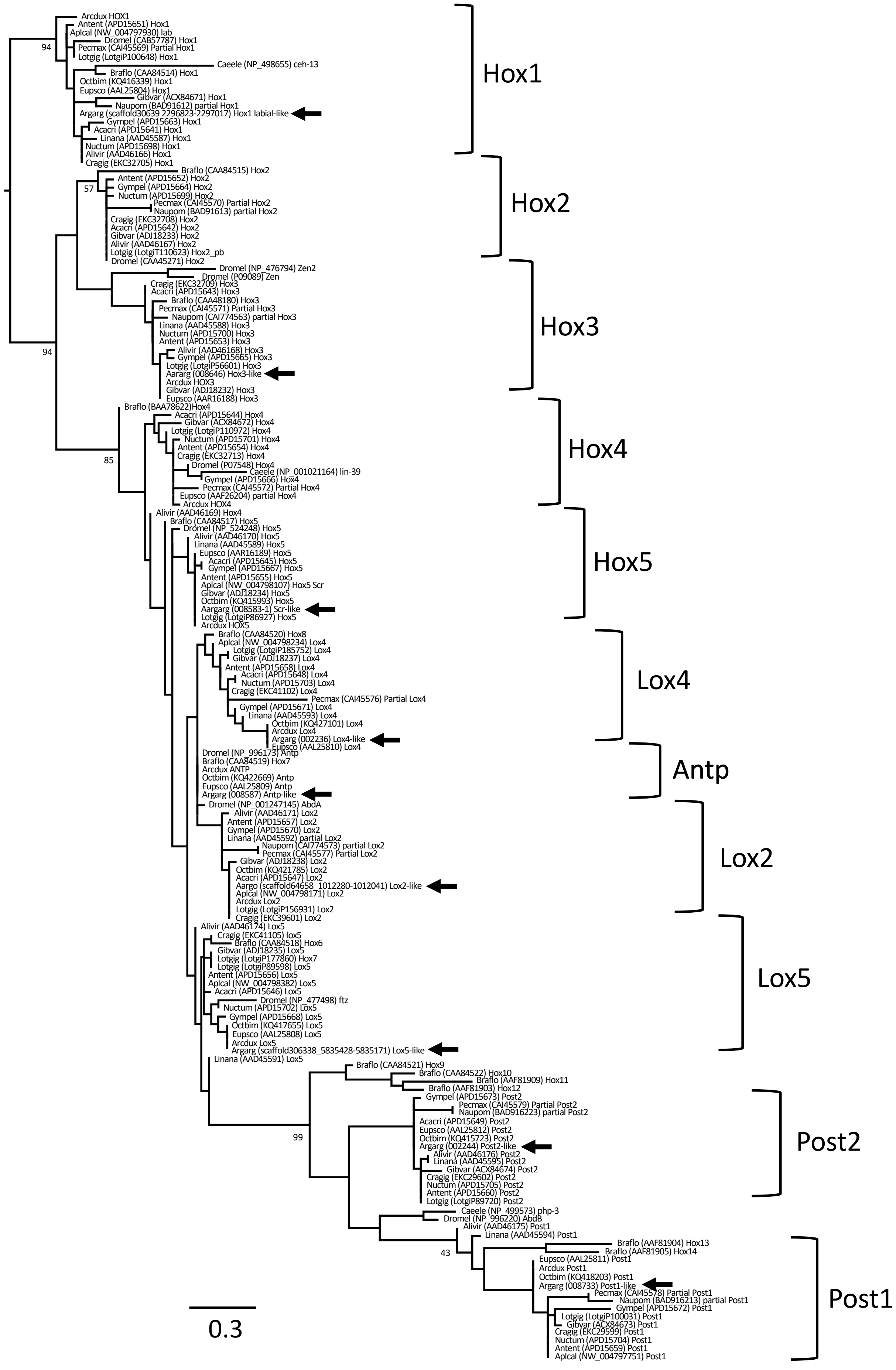


Figure S3 Molecular phylogenetic tree of the Hox genes

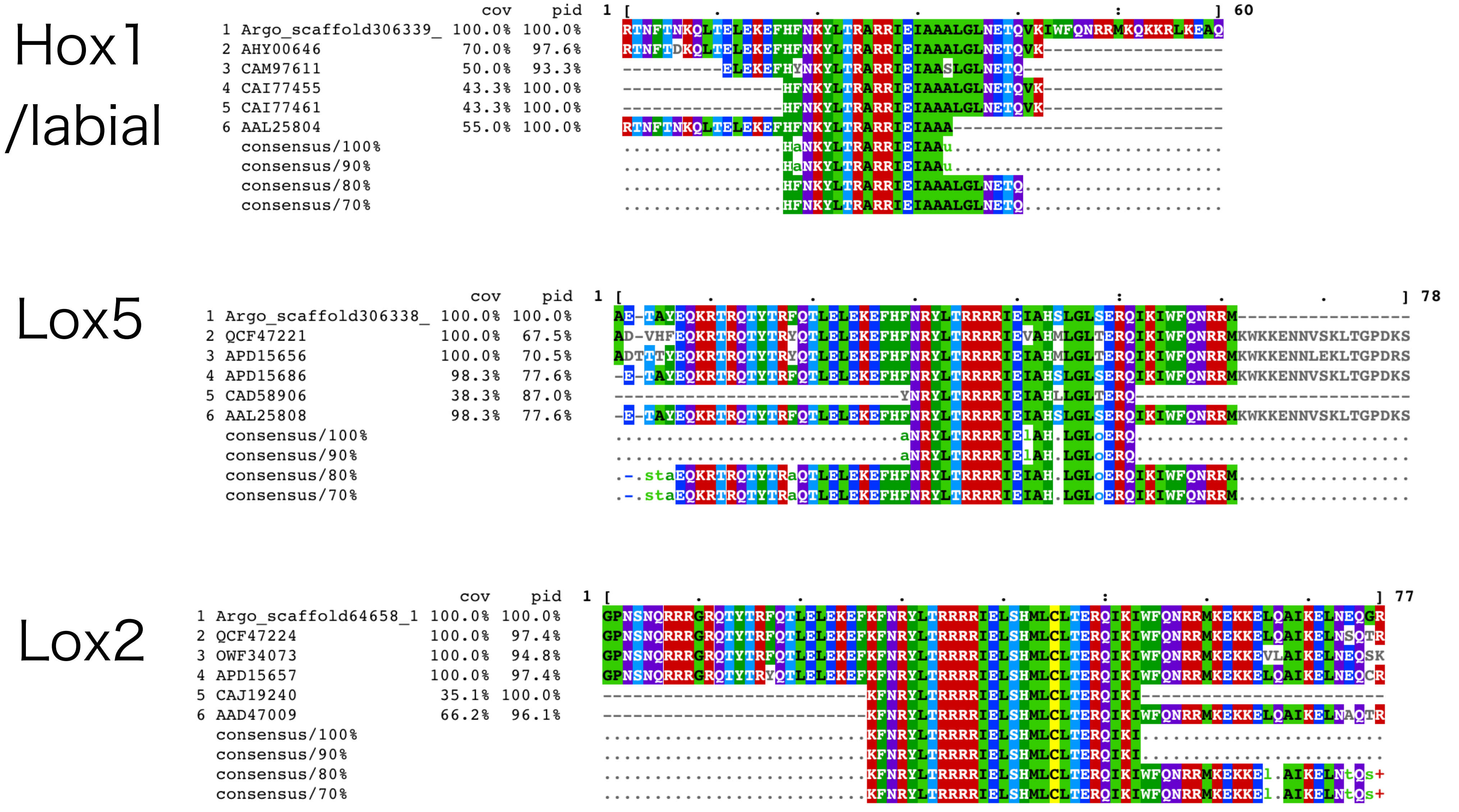


Figure S4 Alignment of non gene model Hox

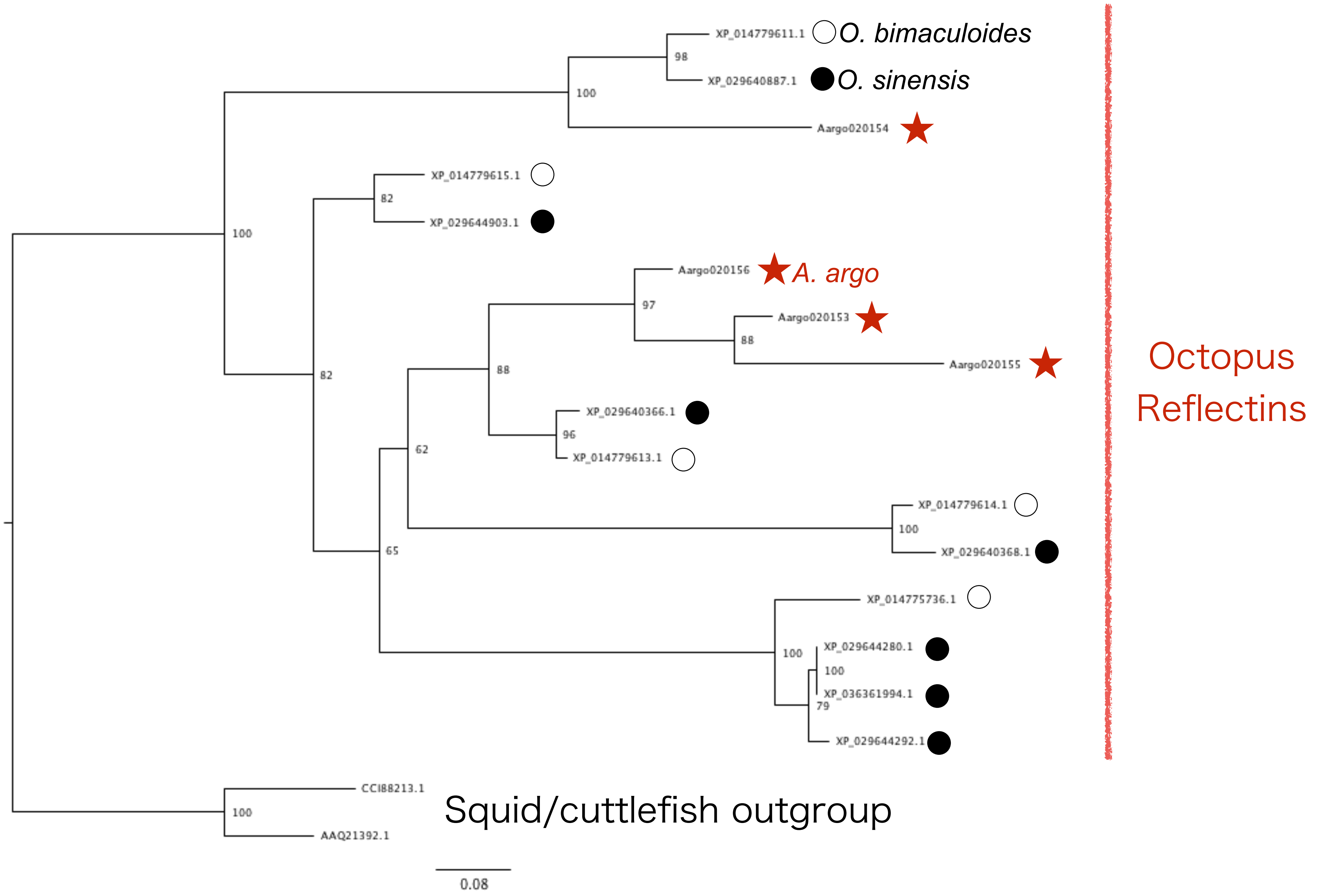


### Figure S5 Reflectin tree

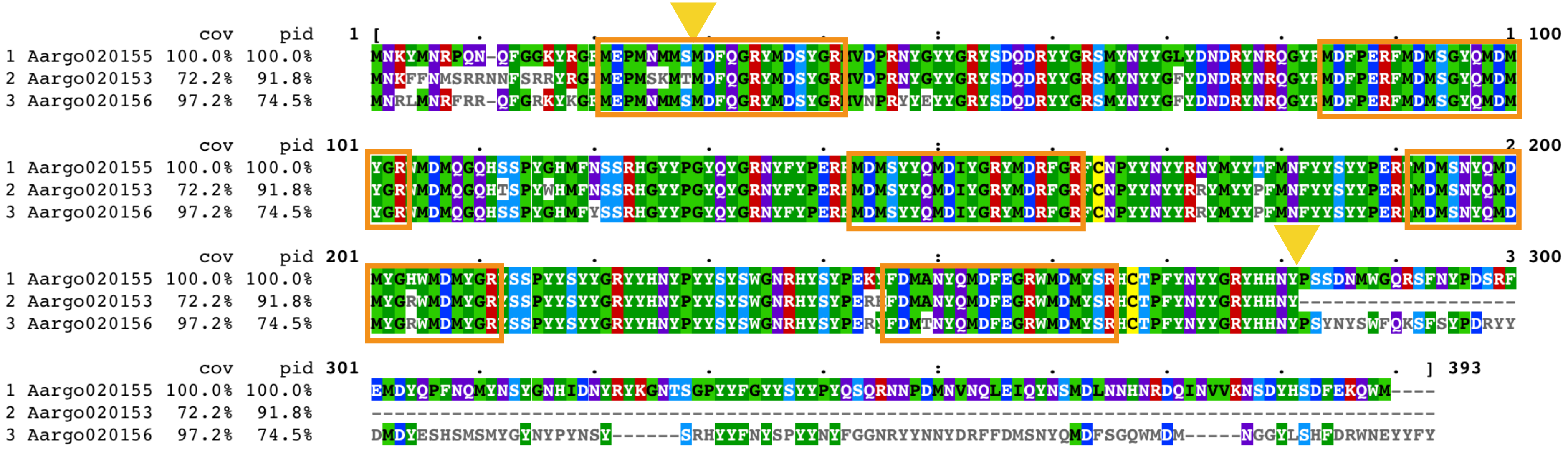

Figure S6 Reflectin alignment with reference to repetitive reflectin motifs

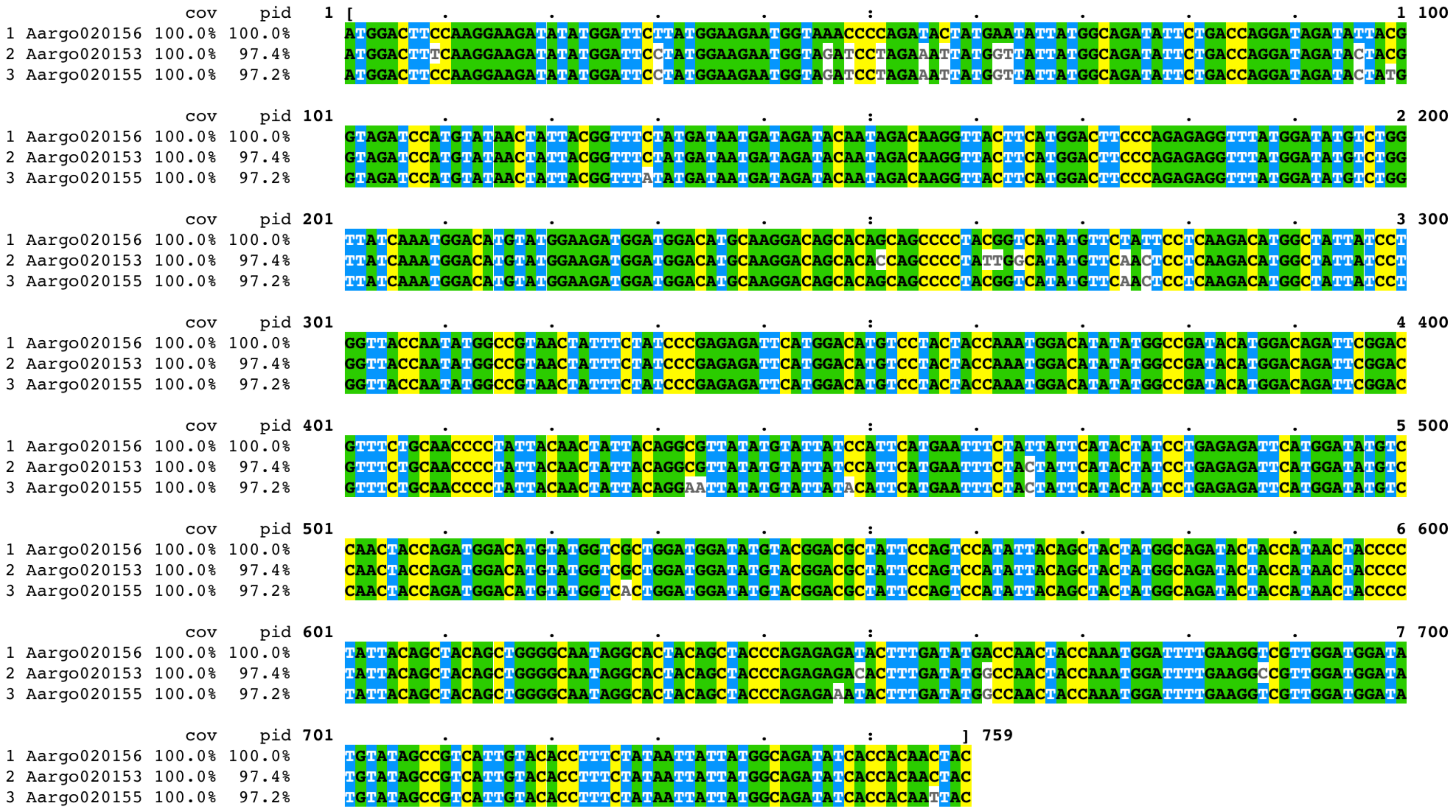


### Figure S7 Reflectin alignment to show gene conversion

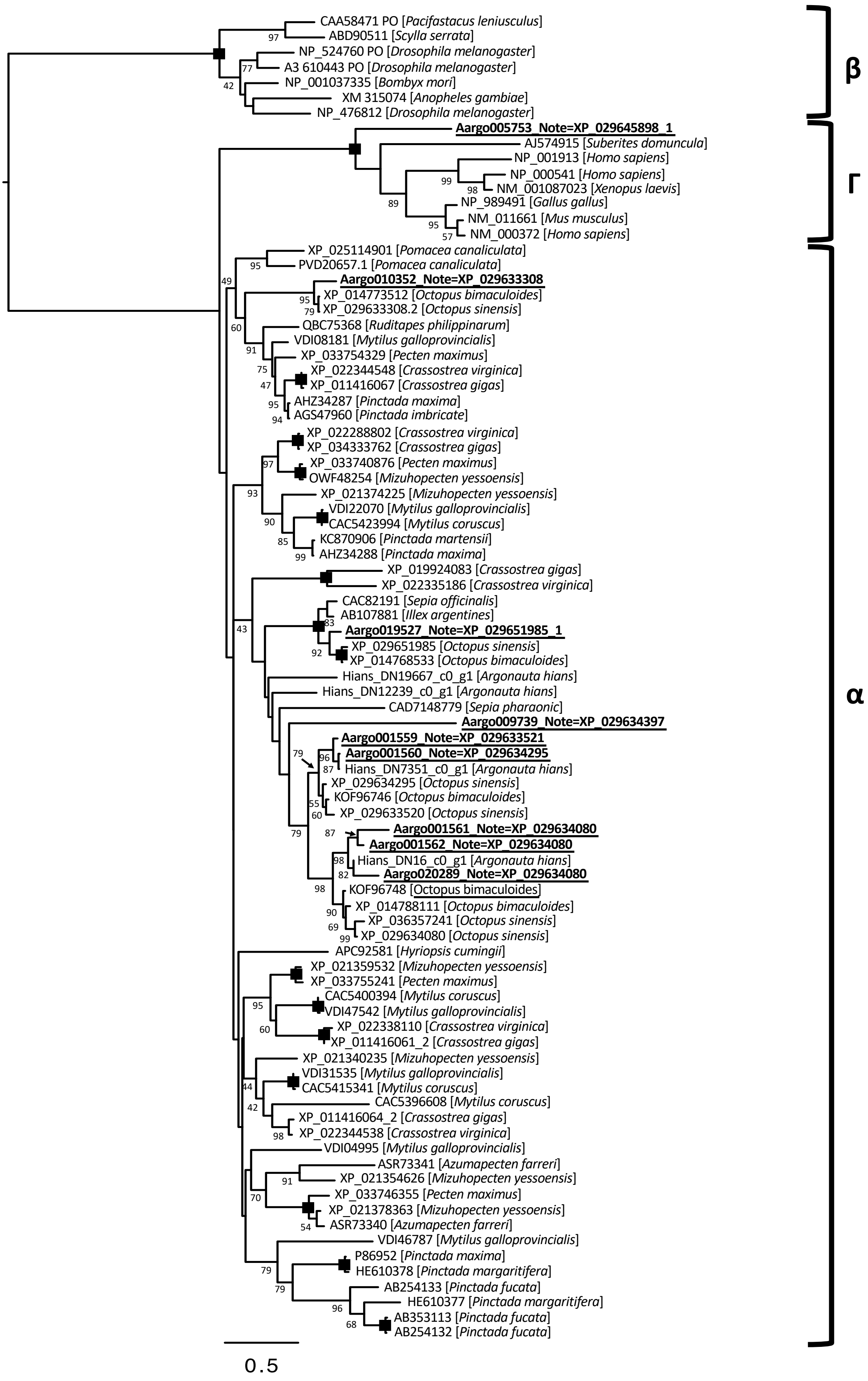


Figure S8 Tyrosinase phylogenetic tree

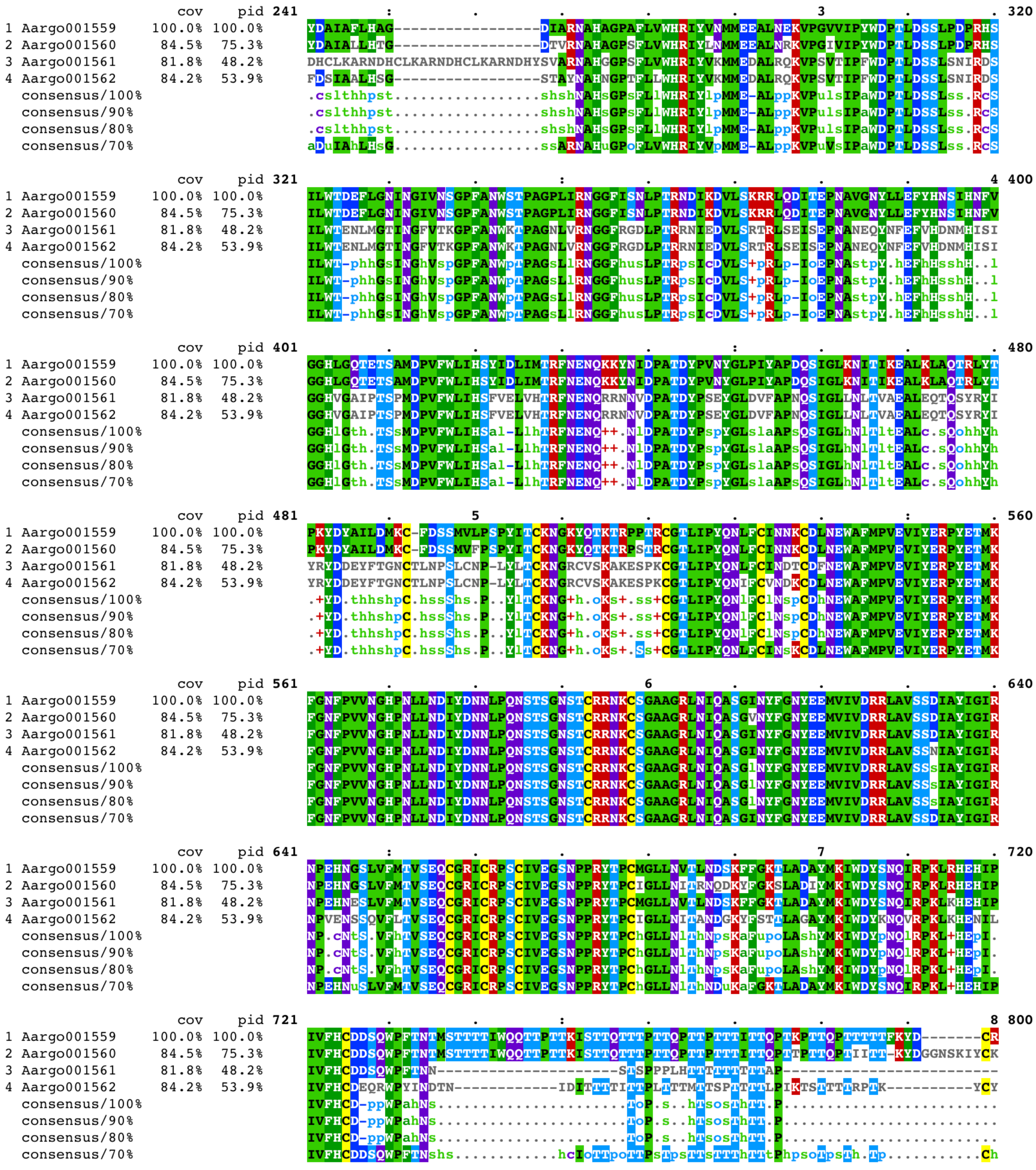

Figure S9 Tyrosinase alignment at amino acid level

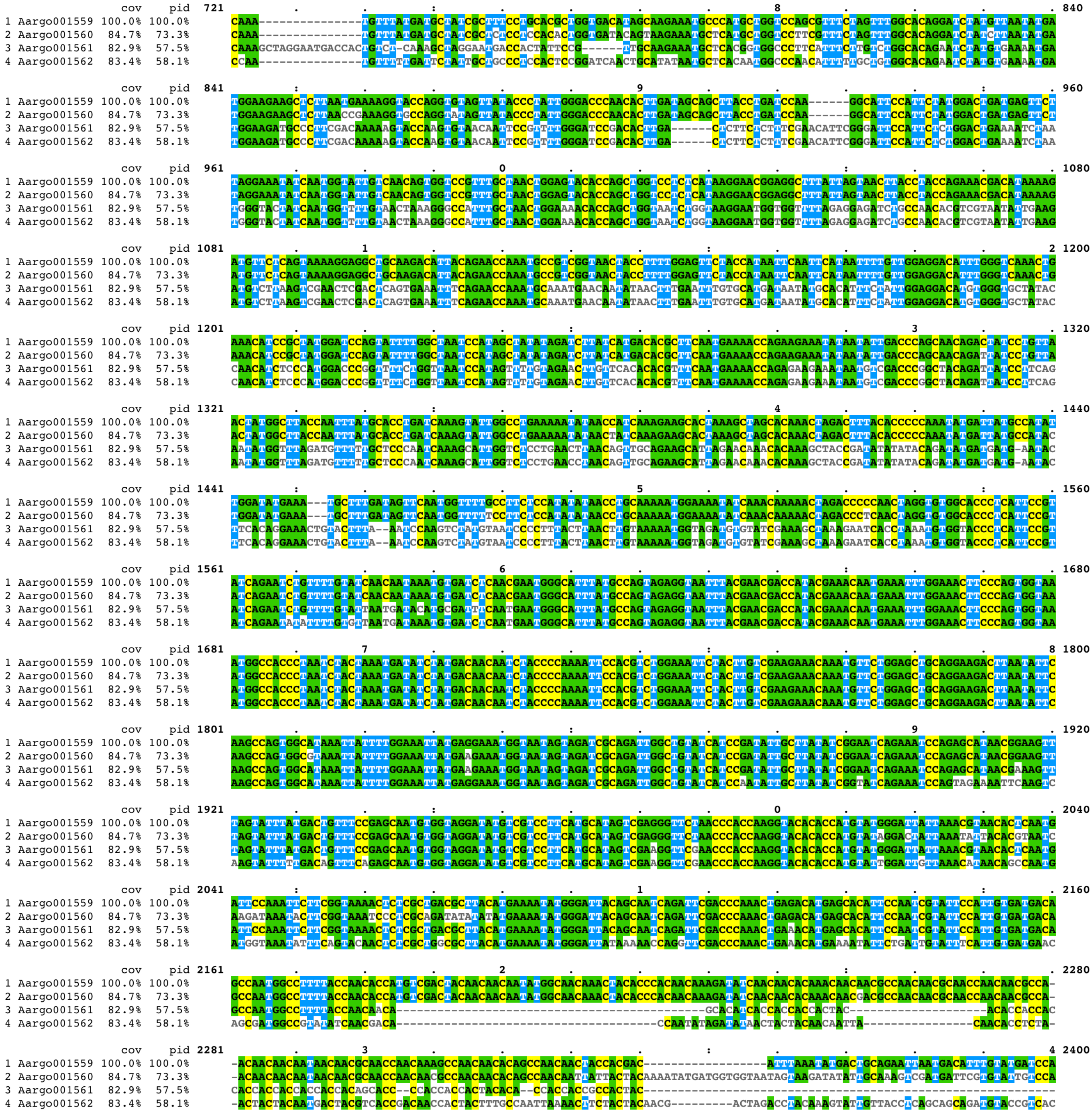

Figure S10 Tyrosinase alignment to show gene conversion

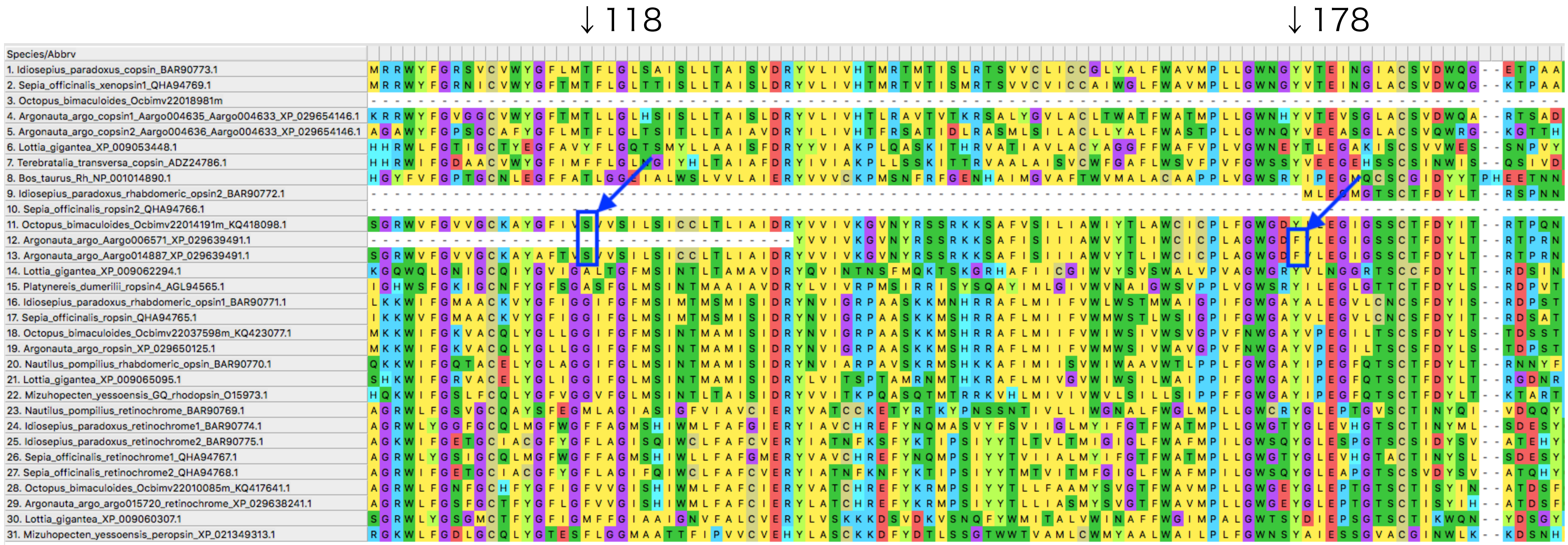


Figure S11 Opsin alignment and unique amino acid chanegs of argonaut based on bovine rhodopsin

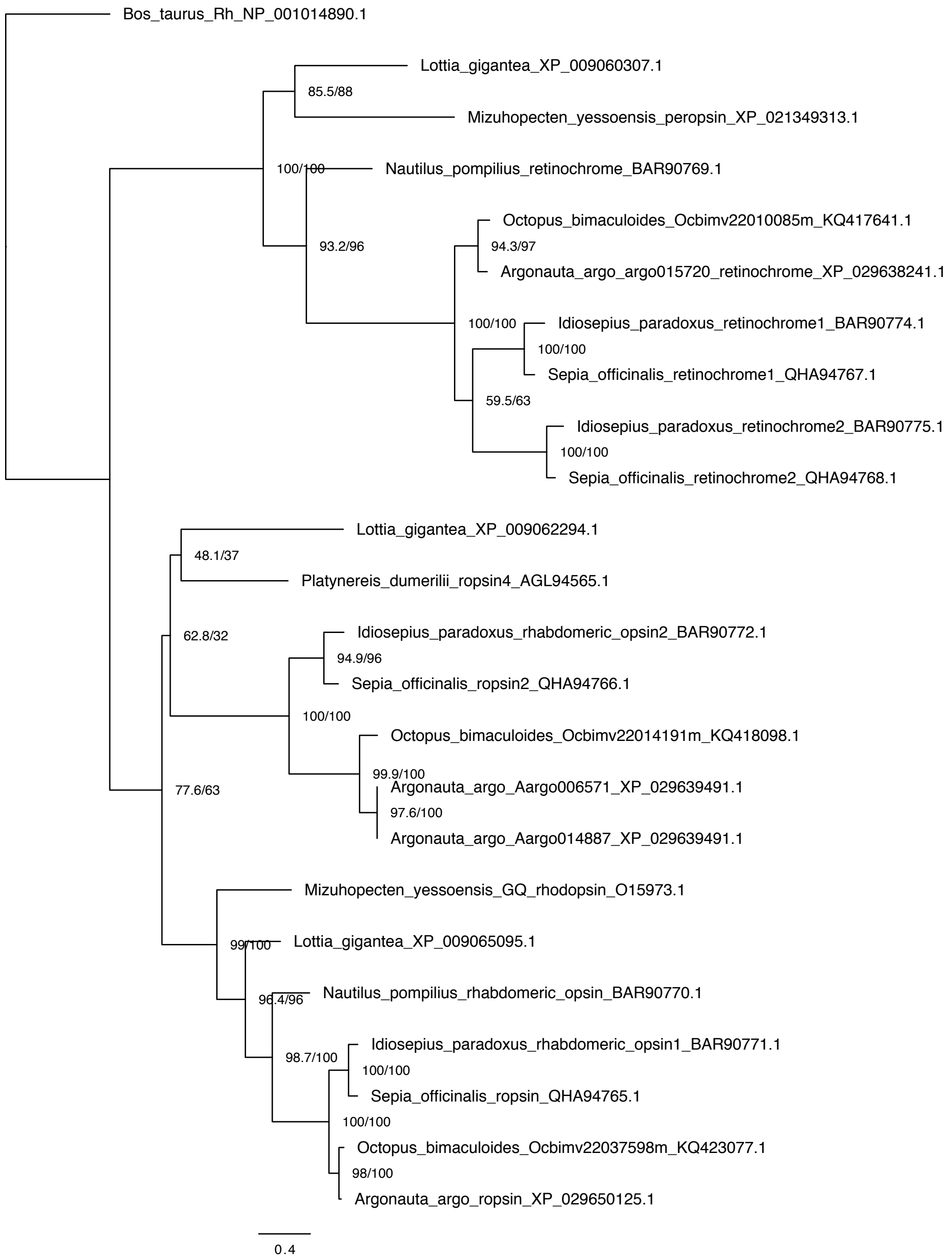


### Figure S12 Phylogenetic tree of RGR and rhabdomeric opsins

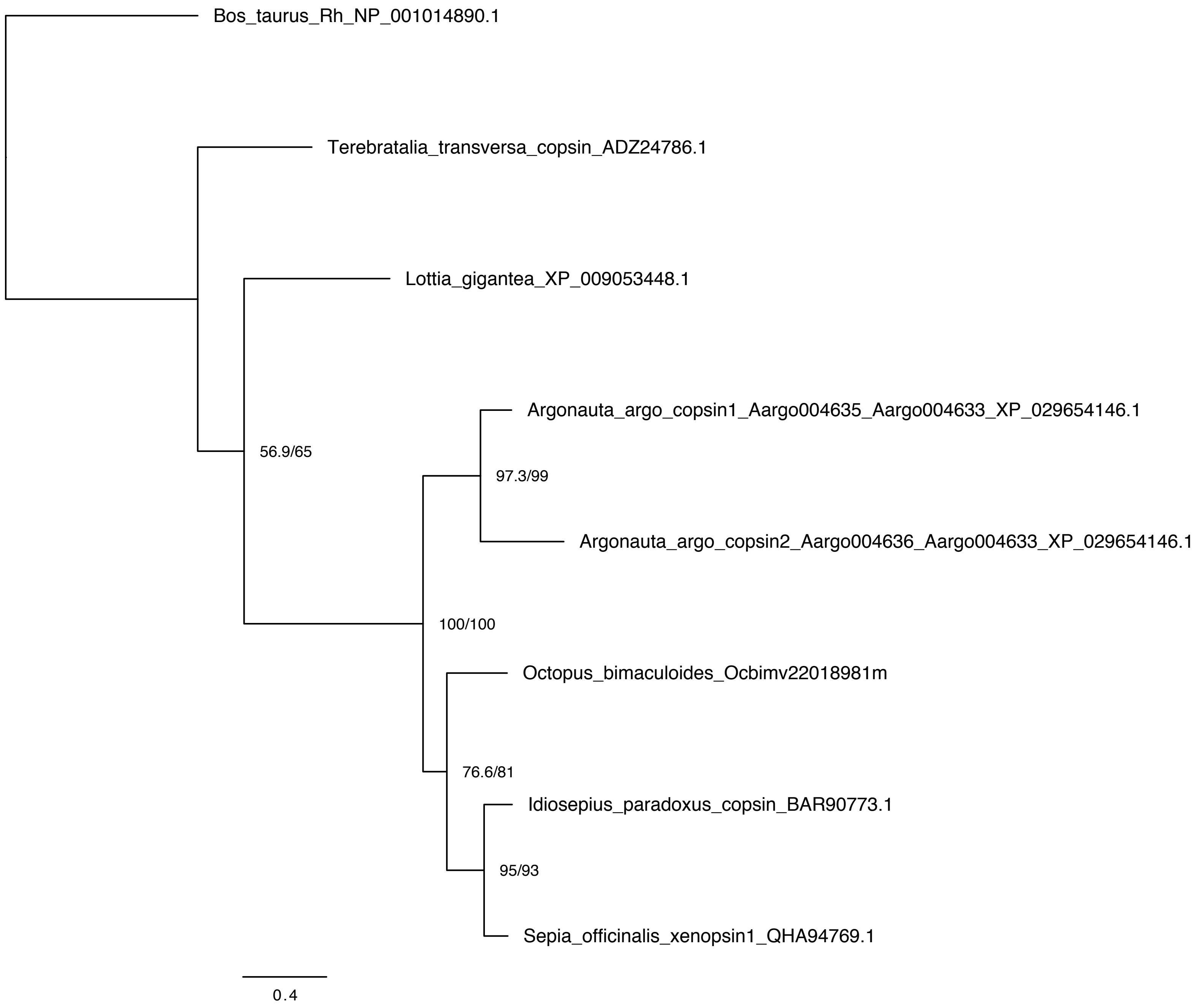


### Figure S13 Xenopsin phylogenetic tree

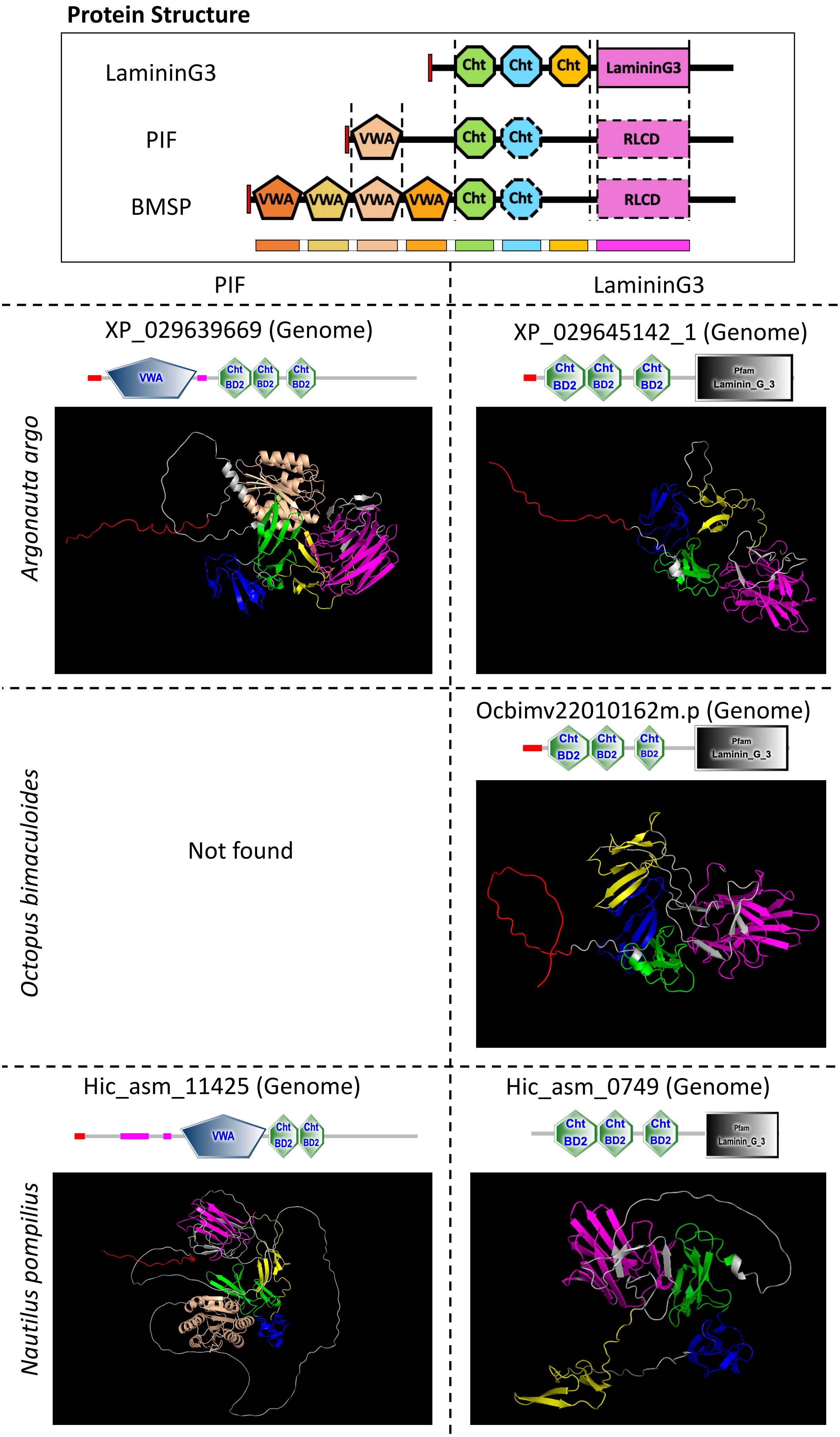


Figure S14 Protein structure of Pif/Pif-like/BMSPs #1

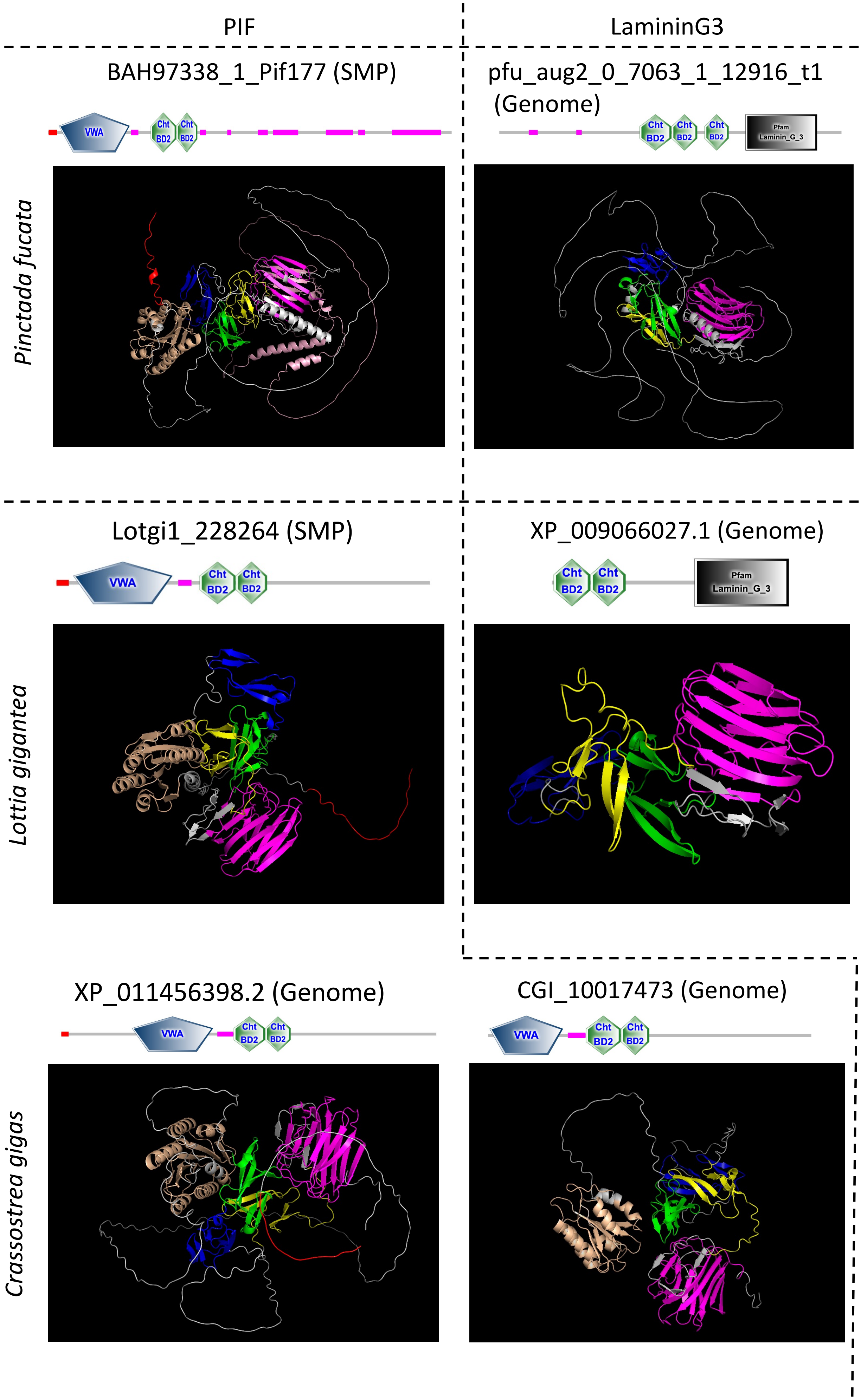


Figure S15 Protein structure of Pif/Pif-like/BMSPs #2

*Mizuhopecten yessoensis*

XP_021370834_1

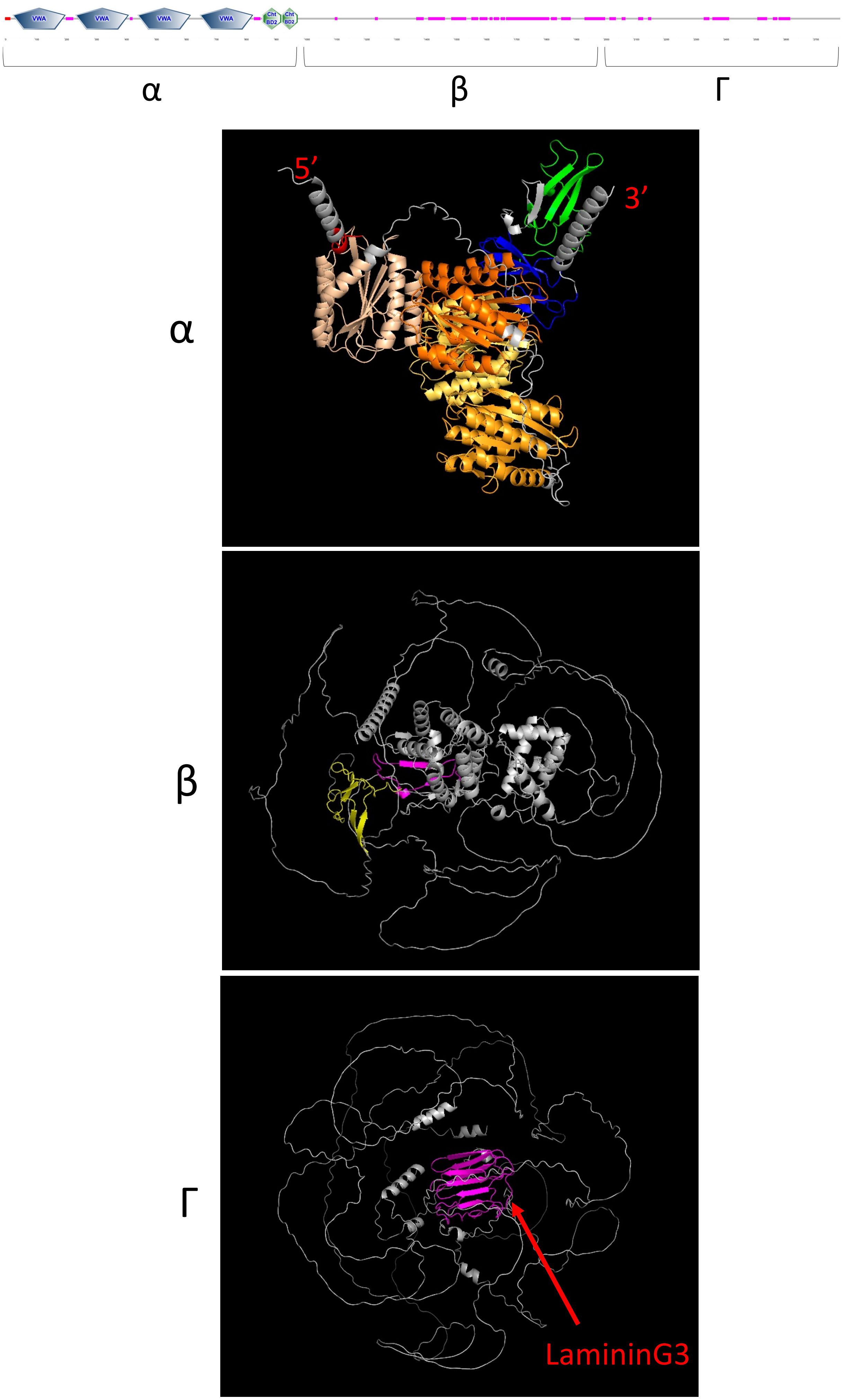


Figure S16 Protein structure of Pif/Pif-like/BMSPs #3
