## Supplementary material for "Gene recruitments and dismissals in argonaut octopus genome provide insights to pelagic lifestyle adaptation and shell-like eggcase reacquisition": Table S

| | Total size (Mbp) | #scaffold | max len (bp) | N50 (bp) | #n50 | gap (%) |
|---|---|---|---|---|---|---|
| This study (Platanus v222) | 1,342.1 | 57,036 | 32,647,630 | 6,178,209 | 56 | 5.30 |
| *Octopus vulgaris* | 2,719.2 | 13,516 | 213,406,131 | **105,892,736** | **10** | **0.03** |
| *Octopus bimaculoides* | 2,305.8 | 48,470 | 4,064,693 | 485,615 | 1,300 | 15.35 |
| *Architeuthis dux* | 2,693.6 | 7,376 | **32,889,981** | 4,852,590 | 160 | 16.76 |
| Crassostrea gigas | 557.4 | **6,433** | 1,964,558 | 402,213 | 399 | 11.82 |
| *Mizuhopecten yessoensis* | 971.6 | 29,088 | 7,498,238 | 827,226 | 334 | 8.23 |

Table S1 Assembly comparison among molluscan genomes

| BUSCO (%) | complete | single | duplicate | fragmented | missing |
|---|---|---|---|---|---|
| This study (Platanus v222) | **91.10** | 88.55 | 2.56 | **3.07** | **5.83** |
| *Octopus vulgaris* | 82.62 | 80.88 | 1.74 | 5.01 | 12.37 |
| *Octopus bimaculoides* | 89.98 | 89.47 | **0.51** | 3.68 | 6.34 |
| *Architeuthis dux* | 88.55 | 87.63 | 0.92 | 3.58 | 7.87 |

Table S2 Assembly comparison based on BUSCO scores (genome mode, metazoa_odb9, n=978)

| *Type* | *Counts* | *Length* (bp) | *Percent* (%) | *Average Length* (bp) | *Relative Abundance* (loci/Mb) |
|---|---|---|---|---|---|
| *Mono* | 32,911 | 444,137 | 1.31 | 13.5 | 25.89 |
| *Di* | 1,424,160 | 40,218,122 | 56.62 | 28.24 | 1,120.52 |
| *Tri* | 729,359 | 18,793,488 | 28.99 | 25.77 | 573.86 |
| *Tetra* | 301,301 | 9,745,400 | 11.98 | 32.34 | 237.06 |
| *Penta* | 18,002 | 433,820 | 0.72 | 24.1 | 14.16 |

Table S3 Microsatellite sequences estimated by Krait

| *GeneID* | *Aargo scaffold ID* | *Hox gene location* | *Best BLAST hit againt Architeuthis genome* | BLAST hit *start* | *end* | *e-value* | *Identity* |
|---|---|---|---|---|---|---|---|
| *Aargo008646* | 306339 | *Hox3* | VCCN01003880.1 | 25694950 | 25695123 | 1.54e-28 | 98.3 |
| *Aargo008543* | 306338 | *Dlx* | VCCN01000387.1 | 16435580 | 16435281 | 1.27e-54 | 94.1 |
| *Aargo008583* | 306338 | *Scr* | VCCN01003880.1 | 24018414 | 24018650 | 2.20e-45 | 100 |
| Aargo008584 | 306338 | | No hit | | | | |
| Aargo008585 | 306338 | | No hit | | | | |
| Aargo008586 | 306338 | | No hit | | | | |
| *Aargo008587* | 306338 | *Antp* | VCCN01003880.1 | 22119788 | 22120258 | 1.75e-45 | 74.1 |
| *Aargo002236* | 64658 | *Lox4* | VCCN01003880.1 | 21387779 | 21388090 | 2.05e-48 | 78.4 |
| Aargo002237 | 64658 | | No hit | | | | |
| Aargo002238 | 64658 | | No hit | | | | |
| Aargo002239 | 64658 | | No hit | | | | |
| Aargo002240 | 64658 | | No hit | | | | |
| Aargo002241 | 64658 | | No hit | | | | |
| Aargo002242 | 64658 | | No hit | | | | |
| Aargo002243 | 64658 | | No hit | | | | |
| *Aargo002244* | 64658 | *Post2* | VCCN01003880.1 | 19080283 | 19080780 | 4.74e-102 | 94.0 |
| *Aargo008733* | 316962 | *Post1* | VCCN01003880.1 | 17446089 | 17445817 | 1.24e-48 | 94.5 |
| *Aargo008772* | 316962 | *En* | VCCN01004895.1 | 231027 | 231302 | 1.35e-46 | 89.1 |

Table S4 Comparison of ORFs in the Hox cluster between blue mussels and giant squid

| Accession | Soluble/InSoluble | Score (Soluble) | Score (inSoluble) | Mass | emPAI (Soluble) | emPAI (inSoluble) | Top hit description |
|---|---|---|---|---|---|---|---|
| Aargo014584 | Soluble/InSoluble | 1265 | 1445 | 18699 | 5.05 | 5.05 | XP_029645183.1(uncharacterized protein LOC115219204) |
| Aargo010955 | Soluble/InSoluble | 3267 | 1685 | 70815 | 3.76 | 2.1 | XP_029648017.1(glypican-6-like) |
| Aargo013239 | Soluble/InSoluble | 2618 | 4552 | 29045 | 3.16 | 6.95 | XP_029645506.1(uncharacterized protein LOC115219476) |
| Aargo009090 | Soluble/InSoluble | 1863 | 5981 | 46847 | 1.25 | 2.11 | XP_029635552.1(kielin/chordin-like protein) |
| Aargo007096 | Soluble | 863 | | 67927 | 0.66 | | XP_029641298.1(kielin/chordin-like protein) |
| Aargo013011 | Soluble/InSoluble | 1088 | 822 | 15665 | 0.61 | 0.61 | no hit |
| Aargo015879 | Soluble/InSoluble | 348 | 395 | 18416 | 0.5 | 1.25 | XP_029639174.1(uncharacterized protein LOC115214207) |
| Aargo018907 | Soluble | 73 | 63 | 9277 | 0.48 | 1.18 | no hit |
| Aargo003237 | Soluble/InSoluble | 39 | 109 | 11360 | 0.38 | 1.64 | XP_029634140.1(histone H4) |
| Aargo008472 | Soluble | 43 | | 27061 | 0.32 | | XP_029657877.1(matrix metalloproteinase-19-like) |
| Aargo019511 | Soluble | 170 | | 56212 | 0.31 | | XP_029657579.1(protein disulfide-isomerase-like) |
| Aargo000068 | Soluble | 94 | | 77205 | 0.28 | | XP_029641298.1(kielin/chordin-like protein) |
| Aargo009082 | Soluble/InSoluble | 572 | 9891 | 137812 | 0.21 | 1.3 | XP_029641298.1(kielin/chordin-like protein) |
| Aargo010041 | Soluble/InSoluble | 108 | 43 | 63890 | 0.2 | 0.06 | XP_029649226.1(70 kDa neurofilament protein-like) |
| Aargo017478 | Soluble | 68 | | 47245 | 0.17 | | XP_029655835.1(collagen alpha-1(XII) chain-like) |
| Aargo003530 | Soluble/InSoluble | 51 | 27 | 23962 | 0.17 | 0.17 | XP_029652506.1(uncharacterized protein LOC115225721) |
| Aargo014807 | Soluble | 38 | | 25623 | 0.16 | | XP_029637330.1(peptidyl-prolyl cis-trans isomerase B-like) |

| | | | | | | | |
|---|---|---|---|---|---|---|---|
| Aargo020077 | Soluble | 45 | | 68646 | 0.12 | | XP_029635321.1(glutathione hydrolase 1 proenzyme-like) |
| Aargo005089 | Soluble/InSoluble | 42 | 35 | 64734 | 0.12 | 0.06 | XP_029636026.1(nucleobindin-2-like) |
| Aargo018523 | Soluble/InSoluble | 34 | 30 | 32307 | 0.12 | 0.12 | XP_029645142.1(uncharacterized protein LOC115219174) |
| Aargo009979 | Soluble/InSoluble | 80 | 92 | 41993 | 0.09 | 0.2 | XP_029636148.1(lysosomal aspartic protease-like isoform X2) |
| Aargo006874 | Soluble/InSoluble | 58 | 74 | 41917 | 0.09 | 0.57 | XP_029639043.1(actin, cytoplasmic) |
| Aargo001902 | Soluble | 54 | | 43454 | 0.09 | | XP_029647804.1(lachesin-like) |
| Aargo019743 | Soluble | 35 | | 41792 | 0.09 | | XP_029643399.1(glyceraldehyde-3-phosphate dehydrogenase-like isoform X1) |
| Aargo012655 | Soluble/InSoluble | 70 | 36 | 50731 | 0.08 | 0.08 | XP_029646382.1(probable peptidylglycine alpha-hydroxylating monooxygenase 1) |
| Aargo005978 | Soluble/InSoluble | 54 | 27 | 101036 | 0.08 | 0.04 | XP_029653787.1(matrix metalloproteinase-2-like) |
| Aargo009281 | Soluble/InSoluble | 50 | 38 | 51291 | 0.08 | 0.08 | XP_029633267.1(elongation factor 1-alpha-like) |
| Aargo005181 | Soluble | 32 | | 52353 | 0.08 | | XP_029636907.1(carbohydrate sulfotransferase 11-like) |
| Aargo009733 | Soluble | 47 | | 60167 | 0.07 | | XP_029640025.1(glypican-3-like) |
| Aargo004396 | Soluble/InSoluble | 42 | 22 | 60268 | 0.07 | 0.07 | XP_029652967.1(vascular non-inflammatory molecule 3-like) |
| Aargo009172 | Soluble | 72 | | 138099 | 0.06 | | XP_029635449.1(Golgi apparatus protein 1-like) |
| Aargo007715 | Soluble | 128 | | 483884 | 0.05 | | no hit |
| Aargo013134 | Soluble | 48 | | 75467 | 0.05 | | XP_029645082.1(uncharacterized protein LOC115219132) |
| Aargo007446 | Soluble | 48 | | 73037 | 0.05 | | XP_029644395.1(endoplasmic reticulum chaperone BiP-like) |
| Aargo003471 | Soluble | 69 | | 184525 | 0.04 | | XP_029656717.1(uncharacterized protein |

| | | | | | | | |
|---|---|---|---|---|---|---|---|
| | | | | | | | LOC115230720, partial) |
| Aargo014038 | Soluble | 48 | | 89243 | 0.04 | | XP_029656725.1(BMP-binding endothelial regulator protein-like) |
| Aargo008197 | Soluble/InSoluble | 47 | 28 | 96408 | 0.04 | 0.04 | XP_029651355.1(calsyntenin-1-like) |
| Aargo017333 | Soluble/InSoluble | 56 | 33 | 116015 | 0.03 | 0.03 | XP_029636473.1(lysosomal alpha-mannosidase-like) |
| Aargo014037 | Soluble | 54 | | 249053 | 0.02 | | XP_029653605.1(uncharacterized protein LOC115226724) |
| Aargo009083 | InSoluble | | 3825 | 60250 | | 1.13 | XP_029641298.1(kielin/chordin-like protein) |
| Aargo003067 | InSoluble | | 41 | 13411 | | 0.32 | XP_029640572.1(histone H2A) |
| Aargo003084 | InSoluble | | 43 | 13546 | | 0.31 | XP_029641645.1(histone H2B, gonadal) |
| Aargo003528 | InSoluble | | 28 | 29098 | | 0.3 | XP_029652588.1(endoplasmic reticulum resident protein 44-like) |
| Aargo002845 | InSoluble | | 22 | 15376 | | 0.27 | XP_029641651.1(histone H3.3) |
| Aargo000067 | InSoluble | | 70 | 88525 | | 0.24 | XP_029641298.1(kielin/chordin-like protein) |
| Aargo002647 | InSoluble | | 35 | 23029 | | 0.18 | XP_029643810.1(ras-related protein Rab-30-like) |
| Aargo013125 | InSoluble | | 29 | 31859 | | 0.13 | XP_029645353.1(uncharacterized protein LOC115219333) |
| Aargo002562 | InSoluble | | 28 | 29878 | | 0.13 | XP_029643815.1(14-3-3 family protein artA-like) |
| Aargo011145 | InSoluble | | 25 | 33590 | | 0.12 | XP_029645177.1(arginase-1-like) |
| Aargo008234 | InSoluble | | 24 | 48249 | | 0.08 | XP_029651017.1(tryptophan--tRNA ligase, cytoplasmic-like) |
| Aargo002156 | InSoluble | | 23 | 56509 | | 0.07 | XP_029640651.1(ATP synthase subunit beta, mitochondrial) |
| Aargo001845 | InSoluble | | 75 | 91175 | | 0.04 | no hit |
| Aargo002777 | InSoluble | | 42 | 90648 | | 0.04 | XP_029653634.1(WD repeat-containing |

| | | | | | |
|---|---|---|---|---|---|
| | | | | | protein on Y chromosome-like) |
| Aargo002913 | InSoluble | 25 | 469668 | 0.01 | XP_029641431.1(spectrin beta chain, non-erythrocytic 1-like isoform X2) |

Table S5 The list of EsMPs based on *Argonauta argo* gene models

| Library types | Read length | Number of reads | Total nucleotides | Total nucleotide after trimming |
|---|---|---|---|---|
| PE600 | 250 | 235M | 117.5Gb | 109.0Gb |
| MP3kb | 100 | 129M | 25.9Gb | 14.7Gb |
| MP6kb | 100 | 130M | 25.9Gb | 16.5Gb |
| MP10kb | 100 | 129M | 25.8Gb | 17.0Gb |
| MP15kb | 100 | 132M | 26.4Gb | 16.8Gb |

Table S6 *A. argo* genome sequencing data

| *Tissues* | Read length | Number of reads |
|---|---|---|
| *heart* | 100 | 26.3M |
| *GH* | 100 | 27.3M |
| eye | 100 | 24.9M |
| *1st* | 101 | 26.0M |
| *2nd* | 101 | 27.4M |
| *mantle* | 101 | 26.8M |

Table S7 *A. argo* transcriptome sequencing data

| | *consensus* | *phased (primary)* |
|---|---|---|
| Number of scaffolds (≧500bp) | 57,036 | 8,338 |
| Total length | 1,342,119,551 bp | 1,105,831,271 bp |
| Longest scaffold | 32,647,630 bp | 13,511,555 bp |
| N50 (scaffold, ≧500bp) | 6,178,209 bp | 1,690,686 bp |
| L50 (#:scaffold, ≧500bp) | 56 | 161 |
| N (bp, %) | 71,142,411 bp (5.3%) | 59,948,776 bp (5.4%) |
| Number of contigs (≧500bp) | 161,761 | 90,477 |
| Total length | 1,266,747,862 bp | 1,044,535,279 bp |
| Longest contig | 443,220 bp | 440,694 bp |
| N50 (contig, ≧500bp) | 22,401 bp | 24,544 bp |
| L50 (#:contig, ≧500bp) | 14,005 | 10,963 |

Table S8 Assembly statistics by *Platanus* v222

| *Prediction* | | *#gene* | *exon/gene* | *#single exon genes* | *Average exon length* (bp) | *Average CDS length* (bp) | *Average intron length* (bp) |
|---|---|---|---|---|---|---|---|
| *RNA-seq* | mapping | 25,952 | 6.44 | 7,777 | 184.16 | 1,186.45 | 3,033.56 |
| | *de novo* | 71,352 | 6.58 | 21,744 | 175.18 | 1,152.55 | 3,145.65 |
| homology | | 121,140 | 11.03 | 10,643 | 143.98 | 1,587.64 | 3,357.91 |
| *ab initio* | Augustus | 25,470 | 4.78 | 7,314 | 221.76 | 1,059.19 | 3,964.98 |
| | SNAP | 80,685 | 4.82 | 5,798 | 113.38 | 546.24 | 4,073.65 |
| Consensus | | 20,293 | 6.9 | 1,393 | 181.51 | 1,252.54 | 3,038.45 |

Table S9 Gene prediction models using custom-made annotation pipeline with transcriptomic data (Inoue et al. 2021)

| BUSCO (%) | complete | single | duplicate | fragmented | missing |
|---|---|---|---|---|---|
| This study (Platanus v222) | **97.03** | 93.56 | 3.48 | **1.43** | **1.53** |
| *Octopus vulgaris* | 97.96 | 96.42 | 1.53 | 0.92 | 1.12 |
| *Octopus bimaculoides* | 94.79 | 93.66 | **1.12** | 2.45 | 2.76 |
| *Architeuthis dux* | 88.45 | 87.12 | 1.33 | 7.87 | 3.68 |

Table S10 Gene model comparison based on BUSCO scores (protein mode, metazoa_odb9, n=978)

| | #gene | Number of exons per gene | #single exon genes | Average exon length (bp) | Average CDS length (bp) | Average intron length (bp) | GT-AG splice (%) |
|---|---|---|---|---|---|---|---|
| This study | 20,293 | 6.90 | 1,393 | 181.51 | 1,252.54 | 3,038.45 | 98.01 |
| *Octopus vulgaris* | 18,183 | 7.93 | 3,121 | 262.50 | 1,542.68 | 6,376.70 | 98.46 |
| *Octopus bimaculoides* | 15,842 | 8.35 | 1,727 | 256.69 | 1,547.02 | 5,280.46 | 98.09 |
| *Architeuthis dux* | 33,406 | 5.08 | 5,290 | 200.48 | 1,015.48 | 3,400.01 | 99.23 |
| *Crassostrea gigas* | 28,402 | 7.91 | 1,358 | 272.60 | 1,483.46 | 923.43 | 98.39 |
| *Mizuhopecten yessoensis* | 24,521 | 8.48 | 1,891 | 345.73 | 1,660.85 | 2,217.62 | 98.86 |

Table S11 Gene model comparison among molluscan genomes
